# Fluidics System for Resolving Concentration-Dependent Effects of Dissolved Gases on Tissue Metabolism

**DOI:** 10.1101/2021.03.07.434330

**Authors:** Varun Kamat, Brian M. Robbings, Seung-Ryoung Jung, John Kelly, James B. Hurley, Kenneth P. Bube, Ian R. Sweet

## Abstract

Oxygen (O_2_) and other dissolved gases such as the gasotransmitters H_2_S, CO and NO affect cell metabolism and function. To evaluate effects of dissolved gases on processes in tissue, we developed a fluidics system that controls dissolved gases while simultaneously measuring parameters of electron transport, metabolism and secretory function. We use pancreatic islets, retina and liver to highlight its ability to assess effects of O_2_ and H_2_S. Protocols aimed at emulating hypoxia-reperfusion conditions resolved a previously unrecognized transient spike in O_2_ consumption rate (OCR) following replenishment of O_2_, and tissue-specific recovery of OCR following hypoxia. The system revealed both inhibitory and stimulatory effects of H_2_S on insulin secretion rate from isolated islets. The unique ability of this new system to quantify metabolic state and cell function in response to precise changes in dissolved gases provides a powerful platform for cell physiologists to study a wide range of disease states.

## INTRODUCTION

### A critical need for instrumentation to study the effect of dissolved gases

Oxygen (O_2_) is a fundamental determinant of cell survival and function in mammalian tissues. In most cells the majority of ATP is generated by oxidative phosphorylation, driven by a series of redox reactions in which O_2_ is the ultimate electron acceptor. Hypoxia is linked to many diseases including stroke, cancer, COVID-19 and diabetic complications. In addition to O_2_, trace gases produced by cells (H_2_S, NO and CO) act as signals to regulate cellular and mitochondrial function^1^. Ischemia-reperfusion injury is a condition common to many disease states and it is thought that during reoxygenation a burst of reactive O_2_ species (ROS) occurs that can damage proteins, lipids and nucleic acids ^2, 3^. Yet despite the scientific and clinical importance of dissolved gases, quantitative methods to measure the real time effects of dissolved gases on intact tissue are not available. Investigators who have studied trace gases and who are characterizing drugs to attain the same benefits ^4^ almost exclusively use aqueous based surrogates/donors of gas. The equivalence of these drugs to the gases they are supposed to mimic has not been tested ^4^. Some investigators have bubbled gas directly into media, but this precludes adding essential protein to the media due to foaming. Thus, there is a strong need to develop technology that enables the study of both abundant and trace dissolved gases. The system we describe here does these analyses both quantitatively and reproducibly.

### A flow culture/assessment system that exposes tissues to precise levels and durations of dissolved gases

We developed a flow culture system according to three fundamental and essential specifications needed to assess effect of gases on tissue: 1. maintain tissue function and viability under continuous flow culture conditions; 2. continuously monitor parameters that reflect intracellular changes in metabolism in real time; 3. precisely control the aqueous and gas phase composition of the media bathing the tissue. Although there are many systems readily available that have some components needed to investigate effects of dissolved gas, none incorporate all three. Commercially available hypoxia chambers (for instance Baker Ruskinn Cell Culture Workstations) control steady state levels of dissolved gas and have been effectively and most commonly used for hypoxia studies ^5, 6^. Microfluidics systems have been developed for establishing cell and tissue models where the 3-dimensional structure and cell-to-cell interactions of native tissue can be recreated, which can be used under steady state gas compositions ^7–9^. However, neither method is designed for implementing, and assessing real time effects of, rapid changes in dissolved gas concentrations The Seahorse flux analyzer measures O_2_ consumption rate (OCR) and extracellular acidification rate (mostly from glycolysis and the TCA cycle) on cell monolayers ^10^ and has been extensively utilized across many fields. However, it is not designed to maintain tissue in physiological buffers or to control dissolved gas levels.

In previous reports we described an earlier version of our flow culture system and demonstrated its ability to maintain a range of tissues (including islets, retina, liver, brain) over hours and days ^11–16^ while continuously assessing metabolic and functional effects of test compounds. This report highlights the incorporation of technology to precisely control both abundant gases (such as O_2_, CO_2_ and N_2_), by using a countercurrent flow device that promotes equilibration between inflow media and premixed gas, and also trace gases, by novel application of permeation tubes. Permeation tubes are commonly used devices for calibration of safety equipment that detect toxic gases such as CO and H_2_S. They consist of liquified gas housed under pressure in a metal jacket, where the gas continuously leaks through a membrane at a steady and accurately calibrated rate. We present in detail the components and operation of the system in the Methods section, and then illustrate the utility of the system to measure the effects of hypoxia followed by reoxygenation on two tissues, pancreatic islets and retina. We also demonstrate how this device can be used to quantify the effects of H_2_S on islet function and liver energetics.

### Assessment of O_2_-sensitive processes: O_2_ consumption rate, reductive state of cytochromes and rate of lactate and pyruvate production

The most direct endpoints with which to assess the acute effects of O_2_ are components of the electron transport chain (ETC), first and foremost OCR. However, measurements of OCR alone cannot identify the mechanisms that are mediating changes in OCR. Under physiological conditions, OCR can increase in response to changes in substrate supply (increased supply of electrons generated from metabolism) and/or demand (as stimulated by ADP ^17^) ^18^ (Fig. 1). Metabolism of fuels is reflected by a proportional change in cytochrome reduction and OCR: as the number of electrons bound to cytochromes increase, OCR increases by mass action (a “push” system). In contrast, increased ATP usage by energy-utilizing cell functions (importantly ion flux and biosynthesis) with a corresponding increase in ADP leads to increased OCR, but without increased cytochrome c reduction (a “pull” system) ^19, 20^. In this way, OCR changes mediated by substrate supply vs. ATP usage can be distinguished, and these fingerprints are informative in understanding mechanisms mediating changes in the ETC induced by H_2_S and O_2_.

**Fig. 1.**
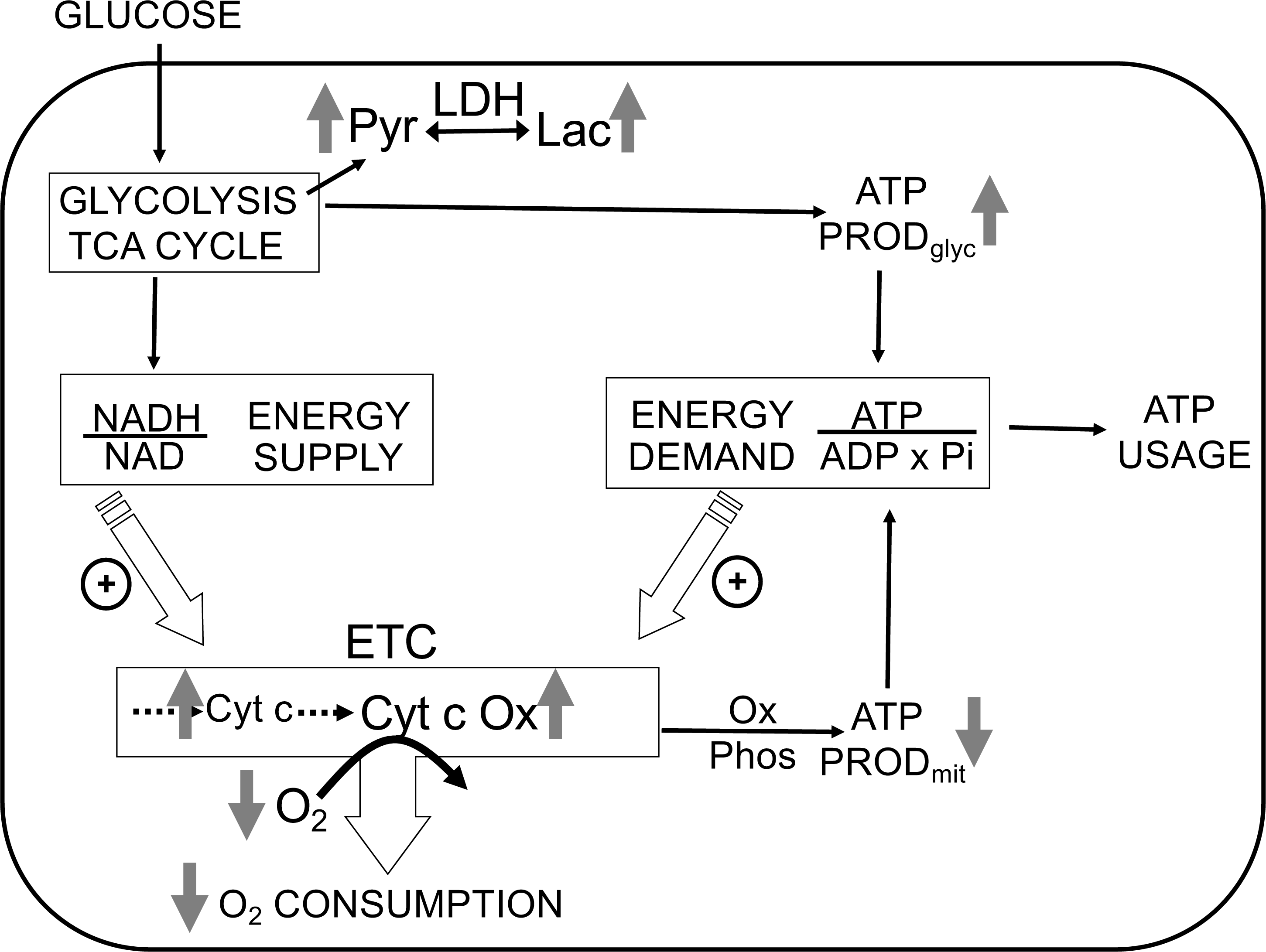
Direct control of OCR: Energy Supply vs Energy Demand vs O_2_. *Schematic depicting three mechanisms mediating OCR.* 1: the supply of reduced electrons generated by metabolism of fuel. 2: the usage of ATP by energy-utilizing cellular function yielding ADP (a major regulator of OCR). 3: the concentration of dissolved O_2_. The concomitant measurement of reduced cytochromes and O_2_ allows for the distinction between the three mechanisms mediating observed changes in OCR. The vertical arrows depict the changes that are acutely affected by changes in O_2_. Low O_2_ leads to increased reductive state of cytochrome c and cytochrome c oxidase, decreased OCR. In some tissue types, the decrease in ATP production by oxidative phosphorylation is compensated by increased ATP generation from glycolysis (the Pasteur effect).

In addition to regulating ETC activity, O_2_ can also influence glycolysis (known as the Pasteur effect ^21^) (Fig. 1). Lactate and pyruvate accumulation and release are determined by relative rates of glycolysis, lactate dehydrogenase (LDH), pyruvate dehydrogenase, mitochondrial and plasma membrane transporters and the redox state of the cytosol. Due to the equilibrium status of LDH, [lactate]/[pyruvate] ratio is proportional to the cytosolic redox state ([NADH]/[NAD]) ^22^, so by measuring both lactate and pyruvate, changes in the rate of lactate production rate due to alterations in glycolytic flux vs. cytosolic redox state can be distinguished. The concomitant measurement of O_2_, OCR, cytochrome c, cytochrome c oxidase, lactate and pyruvate production provide a comprehensive data set to assess the multitude of biochemical and functional effects of O_2_ and other gases.

### Assessment of H_2_S effects on insulin secretion rate (ISR) from isolated pancreatic islets

Like O_2_, H_2_S also interacts directly with the ETC, where it can be both stimulatory and inhibitory. H_2_S can inhibit cytochrome c oxidase ^23^ and it also can donate electrons to cytochrome c ^24^. However, with respect to pancreatic islets, all previous reports have described only inhibition of ISR ^25–30^. Based on the ability of H_2_S to both increase and decrease ETC activity, we chose to demonstrate the technical caliber of our system by testing the hypothesis that H_2_S would both stimulate and inhibit ISR depending on its concentration. Flow culture systems are well-suited to measure changes in ISR in response to changes in perifusate composition ^31^. Secretogogs that affect ISR by isolated islets including glucose, arginine, amino and fatty acids, acetylcholine, GLP-1 as well as sulfonylureas and GLP-1 analogs, also manifest their effects *in vivo*. Therefore, isolated islets are a validated and highly relevant model with which to test our flow culture system. To evaluate the commonly asserted assumption that donors of H_2_S yield the same effects as direct exposure to H_2_S, we also compared the effects of NaHS, a commonly used donor of H_2_S to direct exposure to dissolved H_2_S.

## RESULTS

### Measurement of OCR, reduced cytochrome c and ISR by pancreatic islets in the face of changing inflow O_2_

Ischemia-reperfusion is a stress to tissues that occurs under a range of pathophysiologic conditions, and it is recognized that damage from hypoxia can occur both from the period of decreased energy production and at the time when O_2_ is replenished. Accordingly, we evaluated the ability of our system to measure the recovery of metabolism and function following a period of decreased O_2_ levels. Isolated rat islets were placed into the perifusion chamber and perifused for 90 minutes with Krebs-Ringer Bicarbonate buffer containing 3 mM glucose and equilibrated with 21% O_2_/5% CO_2_/balance N_2_. Changes in OCR, cytochrome c reduction state and ISR, were measured in response to increased glucose (20 mM), decreased O_2_ (by switching to a gas tank supplying the gas equilibration system that contained 3% for 2 hours), and the return of O_2_ to 21% (Fig. 2). To measure OCR, both the inflow and outflow O_2_ concentrations were measured (Fig. 2A), and the data was then processed by convolution techniques described in the Methods section. After transformation of the inflow O_2_ using eq. 4 and the transfer function generated with data obtained in the presence of potassium cyanide (KCN) to account for the delay and dispersion of the perifusion chamber, OCR was calculated from eq. 5. As expected, OCR increased with increased glucose concentration, and decreased with lower O_2_ tension (Fig. 2B). Unexpectedly and notably, there was a transient spike of OCR when O_2_ in the media was restored. OCR then approached a steady state that was about 55% of the pre-hypoxic level of OCR. One limitation of the system arises when defining the O_2_ levels that tissue is exposed to, when in fact there is a gradient from the inflow to the outflow. This uncertainty can be minimized by increasing flow rate thereby decreasing the difference between inflow and outflow concentrations; however, as the difference gets smaller, the resolution of the method decreases.

**Fig. 2.**
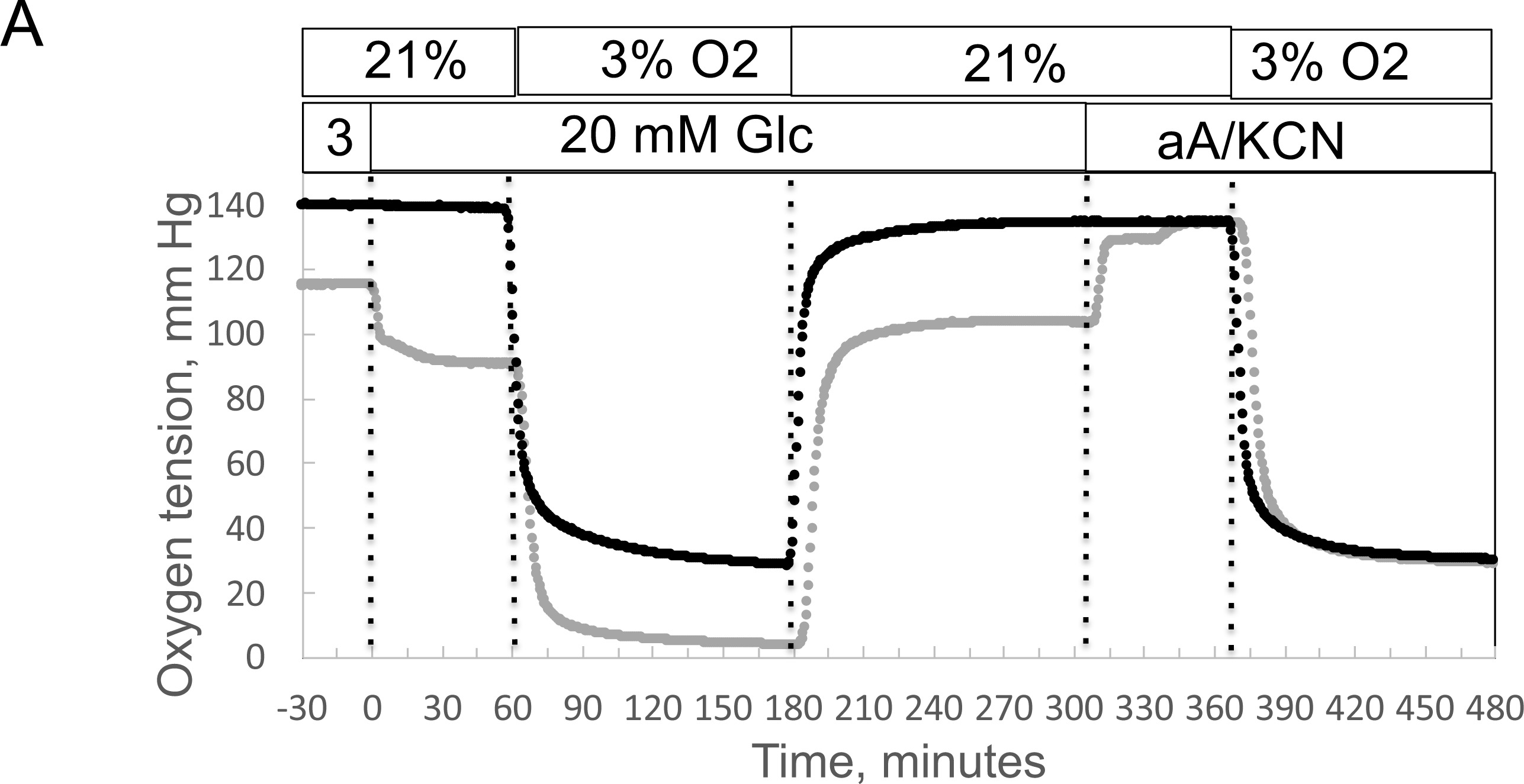
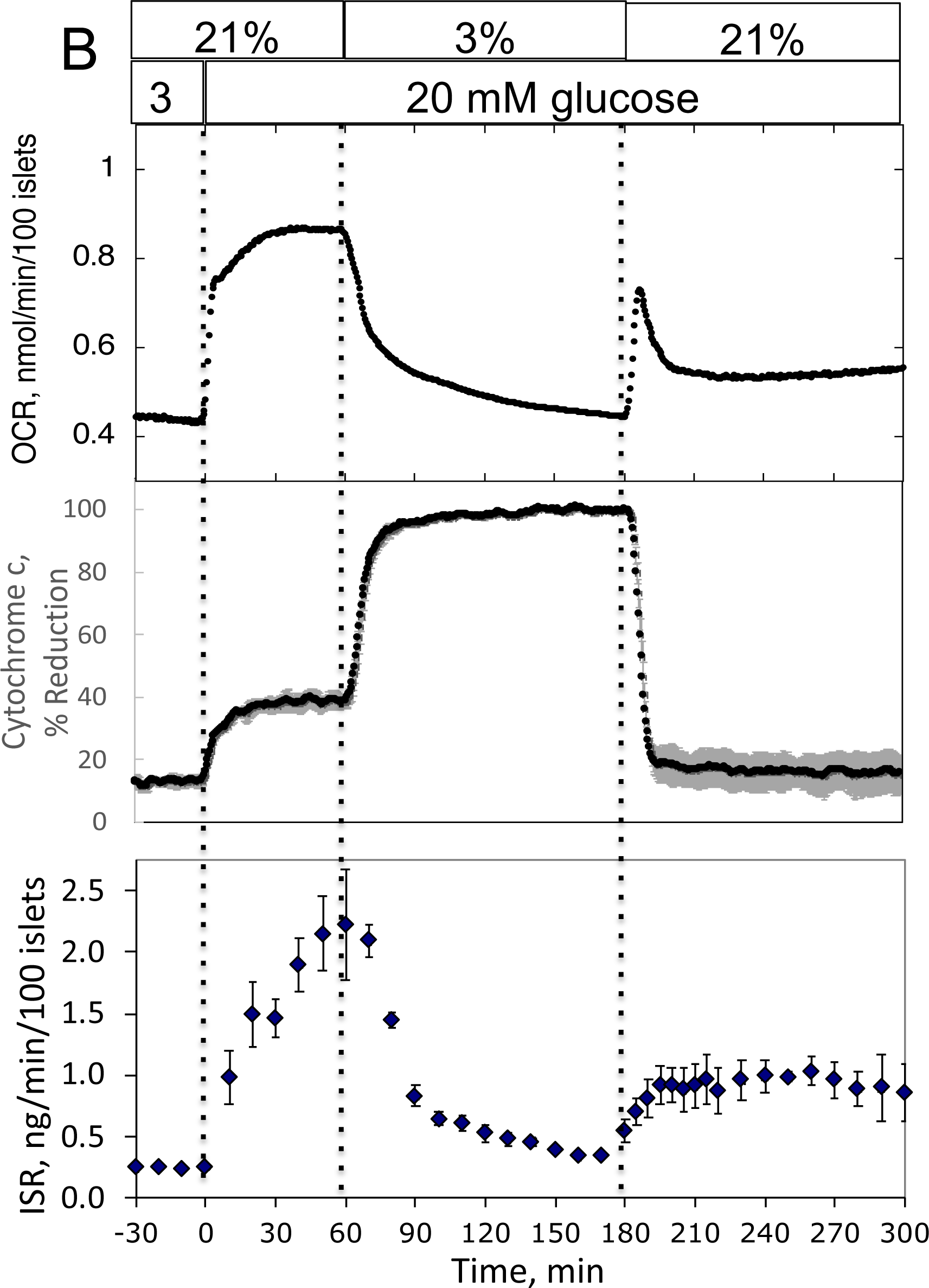
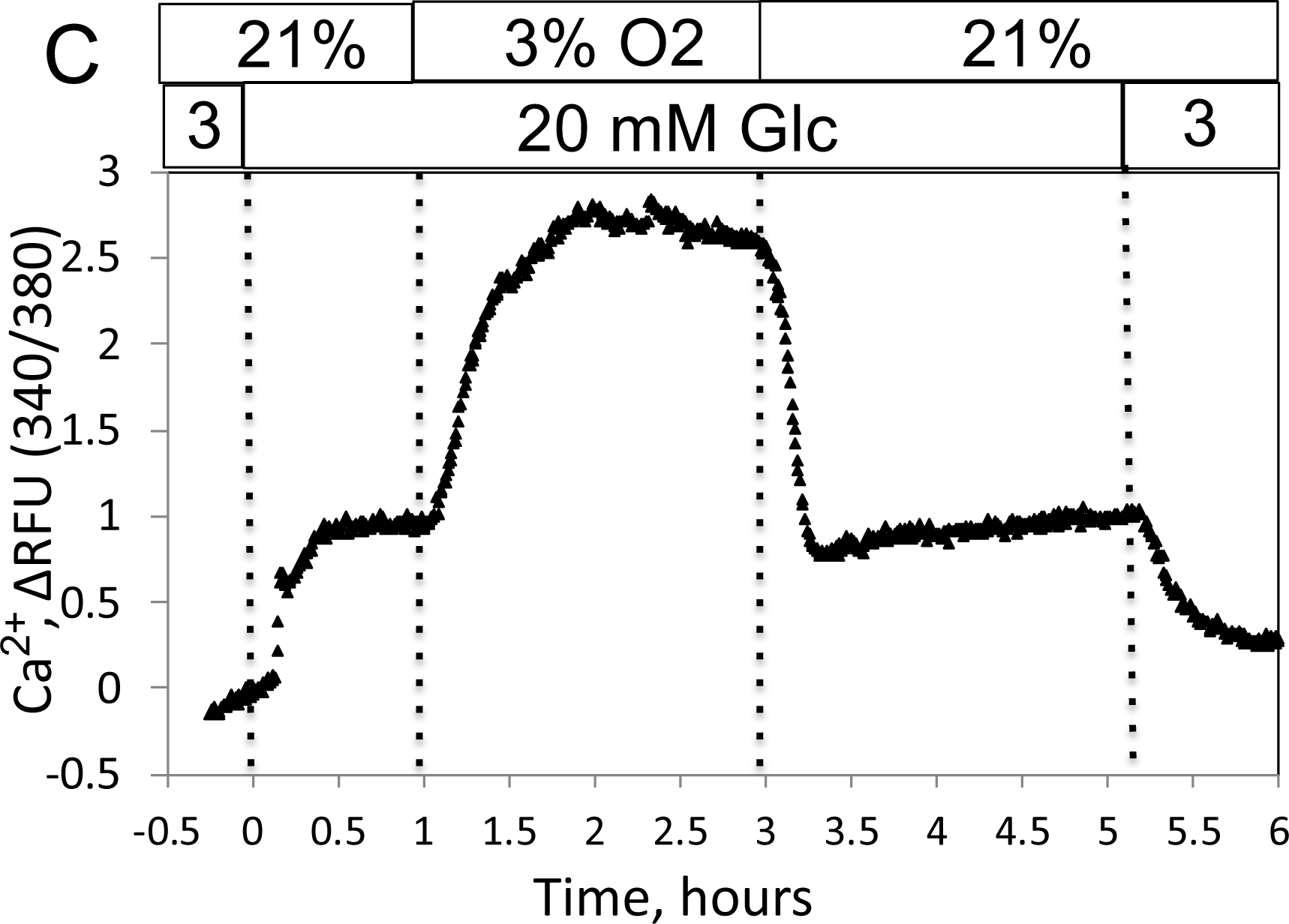
Effect of glucose and low O_2_ on transient and steady state ETC, and ISR and intracellular Ca^2+^ in pancreatic islets. Islets (900/channel) were handpicked with a P200 pipet and after mixing with Cytodex beads (1 uL/10 islets) loaded into the perifusion chamber, and the flow rate was set to 150 uL/min with Krebs-Ringer Bicarbonate buffer containing 3 mM glucose for 90 min. At time = 0, glucose concentration was raised to 20 mM for 45 min; subsequently, O_2_ was decreased to 3% for 2 hours, and then returned to 21%. **A.** The protocol generated inflow and outflow O_2_ profiles such as shown. Following the completion of the protocol, 12 μg/ml antimycin A (aA) was added for 25 minutes, and then 3 mM KCN, and the hypoxia protocol was repeated while islet respiration was suppressed in order to characterize delay and dispersion due to the separation in space of inflow and outflow sensors. **B.** Calculated values of OCR (representative data from an n of 3 (average recovery = 0.55 +/− 0.07), reduced cytochrome c (n =2), and ISR (n =2) were plotted as described in the Methods section. **C.** In a separate illustrative experiment, intracellular Ca^2+^ in islets was imaged and quantified using the same protocol except glucose was also decreased back to 3 mM glucose at the end of the experiment.

The reduction state of cytochrome c was concomitantly measured with OCR (Fig. 2B). Previous reports described an equilibrium with respect to the flow of electrons between NADH and cytochrome c ^32, 33^, and their reductive state represents a balance between the supply of electrons generated by metabolism of fuels and use of electrons to drive proton translocation and ATP production. Consistent with these scenarios, glucose provided more reducing power to drive cytochrome c to its reduced state (in parallel with OCR), whereas hypoxia favored the reduced state of cytochrome c by slowing its oxidation. Following the return to 21% O_2_, cytochrome c reduction reached a steady state of 42% of the pre-hypoxic levels, consistent with incomplete recovery of OCR. A particularly powerful feature of the systems approach is realized when tissue function can be measured concomitantly with measures of ETC. Fractions were collected during the protocol that were later assayed for insulin (Fig. 2B). Stimulation of ISR by glucose in the presence of 21% O_2_ was suppressed in low O_2_, and ISR recovered to 44% of its original level of stimulation after reoxygenation.

### Measurement of calcium (Ca^2+^) in response to decreased O_2_ by islets

As fluorescence imaging is a powerful and widely used modality to assess many intracellular signals including but not limited to Ca^2+^ ^34^, NADH^35, 36^, mitochondrial membrane potential^37^, ATP ^38^, and ROS^11^ in islets, we demonstrated the control of dissolved gas for this modality. We measured the effect of glucose and hypoxia on islet intracellular Ca^2+^ with a protocol similar to the one we used for OCR except that glucose was lowered back to 3 mM at the end of the experiment (Fig. 2C). As expected, intra-islet Ca^2+^ increased in response to the increase in glucose. Subsequently, in response to a decrease in O_2_, Ca^2+^ fluorescence rose 3-fold, presumably reflecting a loss of energy-dependent pumping of Ca^2+^ out of the cells. Notably, in contrast to OCR and ISR parameters, Ca^2+^ recovered fully to pre-hypoxic levels when O_2_ was returned to normal levels.

### Measurement of lactate/pyruvate production and release by perifused INS-1 832/13 cells in response to changes in O_2_

To track shifts between glycolytic and mitochondrial energy generation in real time, fractions collected from the outflow were assayed for lactate and pyruvate. We predicted that extracellular ratios of these two analytes reflect intracellular regulation of these two compounds. To evaluate this, we measured the response to inhibitors of LDH and mitochondrial transport of pyruvate in INS-1 832/13 cells (henceforth referred to as INS-1 cells). INS-1 cells were used instead of islets because most of the pyruvate made in islets is transported into mitochondria ^39^ since they don’t have significant capacity for plasma membrane transport of lactate or pyruvate ^40^. Oxamate, an inhibitor of LDH, rapidly and completely suppressed lactate release from cells (Fig. 3A), showing the tight relation between production of lactate from LDH and appearance of lactate in the outflow. Somewhat surprisingly, pyruvate did not increase. However, OCR increased suggesting that the decrease in flux from pyruvate to lactate was counterbalanced by an increase in flux of pyruvate into the mitochondria. Blocking transport of pyruvate into mitochondria with zaprinast (a blocker of the mitochondrial pyruvate carrier (MPC) ^41^ led to a rapid increase in pyruvate release from cells (Fig. 3B) and diminished OCR. Why lactate production decreased isn’t clear from the data but could be explained by a decrease in the cytosolic redox state (NADH/NAD) by the increased activity of the malate/aspartate shuttle. Thus, the rapidity of the changes in lactate and pyruvate production in response to changes in LDH and MPC indicates that membrane transport is fast, and at least in this cell line, extracellular lactate and pyruvate will reflect changes in intracellular events controlling lactate and pyruvate with a time delay of no more than a few minutes. To ensure that this is the case, these experiments would have to be done in whatever tissue is being investigated.

**Fig. 3.**
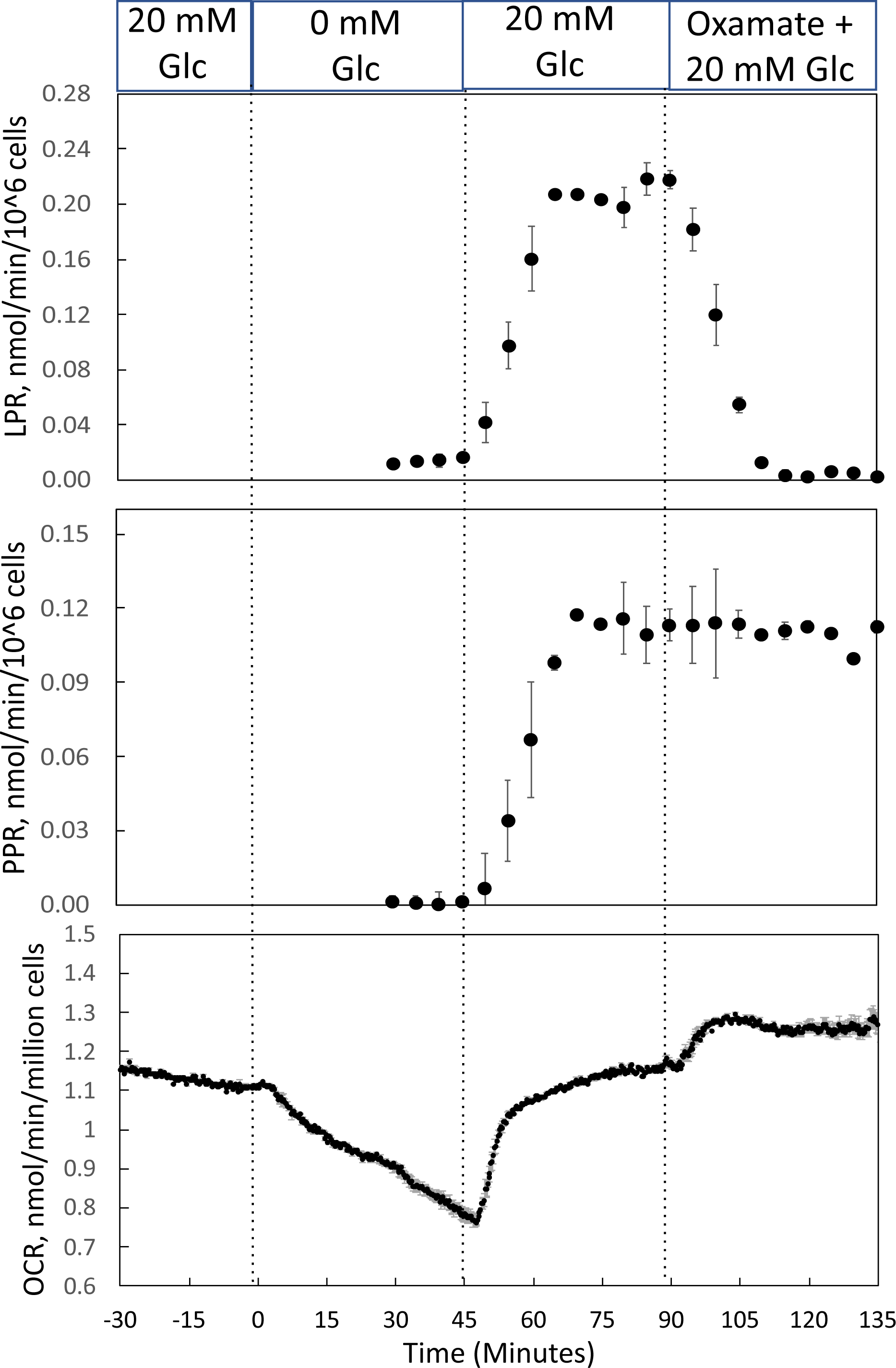
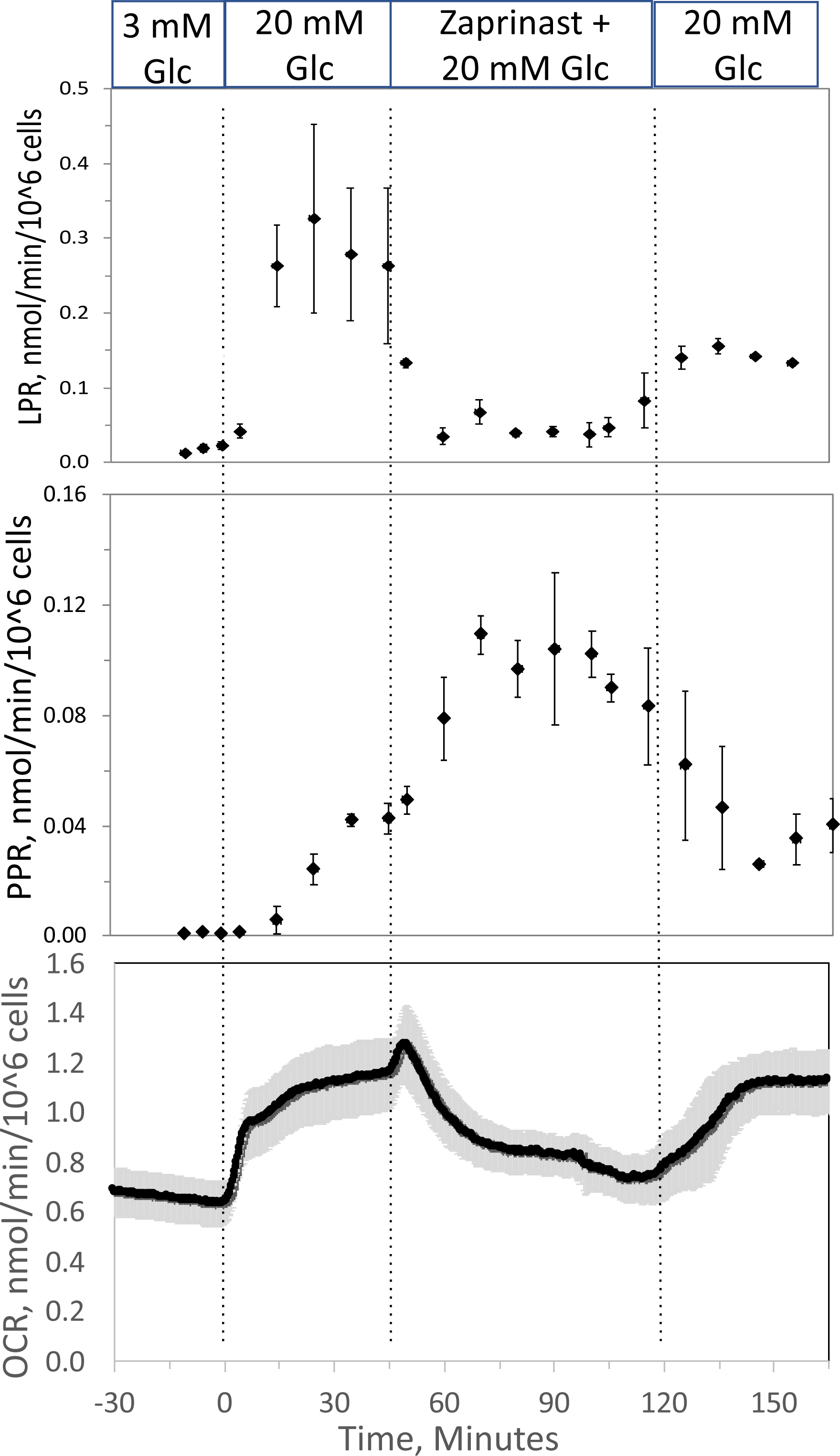
Effect of glucose and inhibitors of LDH and MPC on OCR, lactate and pyruvate production rate by INS-1 832/13 cells. **A.** The effects of glucose and then oxamate (50 mM, an inhibitor of LDH) on OCR, lactate production rate and pyruvate production rate at the times indicated in the figure were measured. **B.** The effects of glucose and then zaprinast (200 μM, an inhibitor of MPC) on OCR, lactate production rate and pyruvate production rate. Data for both plots are the average +/− SE, n =2).

### Measurement of OCR, cytochrome c, lactate and pyruvate by perifused retina before and after a period of hypoxia

In order to compare results with islets to those obtained by a tissue that is less sensitive to hypoxia, experiments were carried out on isolated retina, a tissue that normally resides at low O_2_ ^16^. Similar to the measurement in islets, the inflow and outflow O_2_ were measured (Fig. 4A), and OCR was calculated after convolution of the inflow data. OCR decreased at low O_2_, and then manifested a transient spike in response to reoxygenation (Fig. 4B). However, in contrast to islets, OCR by retina then approached a much higher recovery (a steady state of 83% of the pre-hypoxic rate). Reduced cytochrome c increased to maximal levels during 1% O_2_ and stayed at this level throughout the 2 hours of low O_2_ (Fig. 4B). Consistent with OCR data, upon return of O_2_ to 21% cytochrome c reduction recovered to 79% of pre-hypoxic levels. Comparing post-hypoxia levels of OCR in retina vs. islets (Fig. 4C), islets did not recover from hypoxia as well as retina suggesting that the approach can be used to assess sensitivity to the stress of ischemia-reperfusion conditions. Note that due to the delay in time it took for the inflow perfusate to reach a new equilibrium, the inflow O_2_ did not reach equilibrium levels with the O_2_ from the supply gas tank. The levels of O_2_ in the outflow are dependent upon the inflow O_2_, the flow rate and the OCR of the tissue in the chamber. In order to match the levels of O2 that each tissue was exposed to, the supply gas tanks used for the hypoxia phase of the experiments were selected to generate similar outflow O_2_ levels for the retina (1% yielded an outflow of 6.5 mm Hg) vs. islets (3% yielded 5.5 mm Hg).

**Fig. 4.**
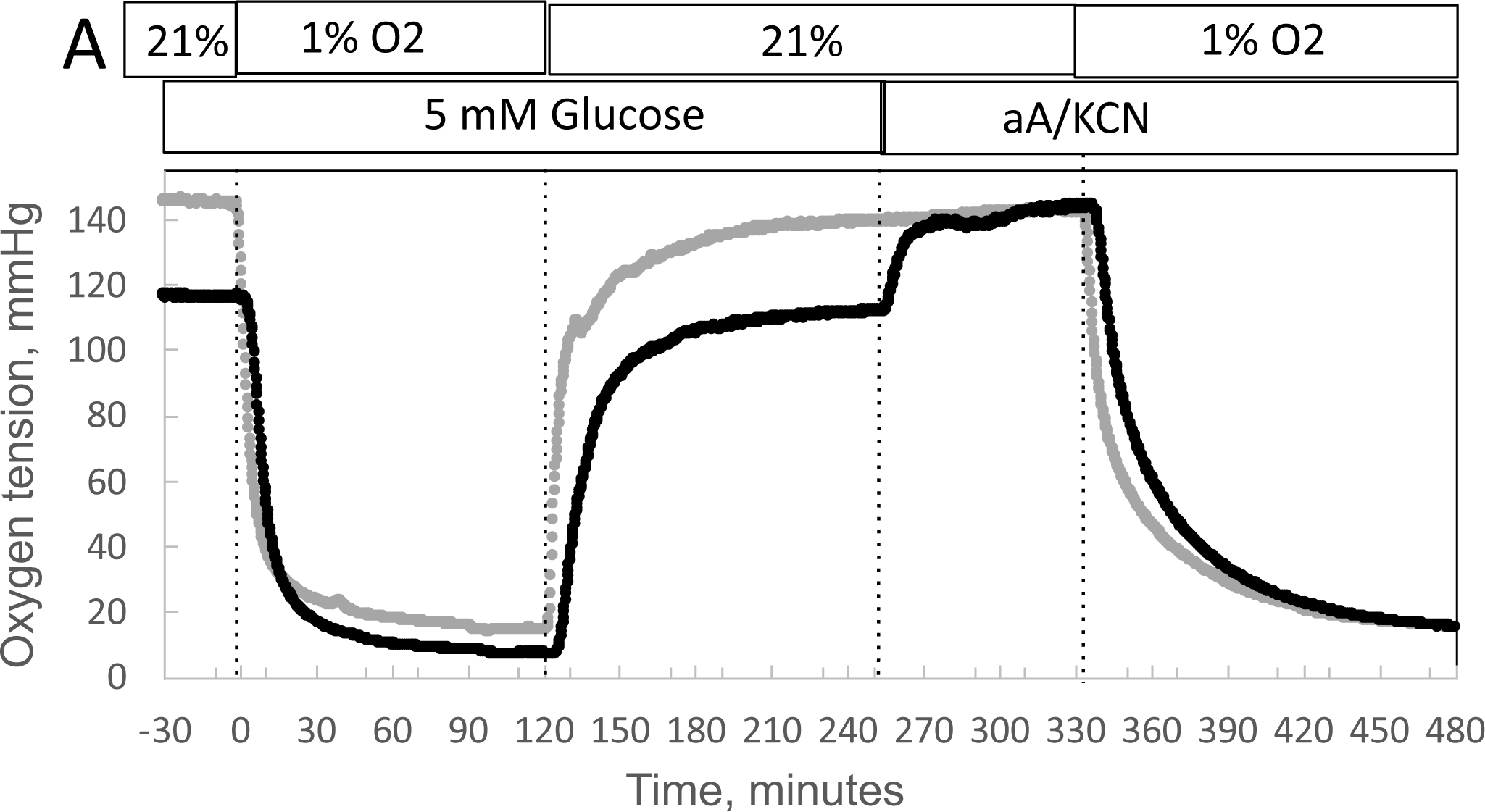
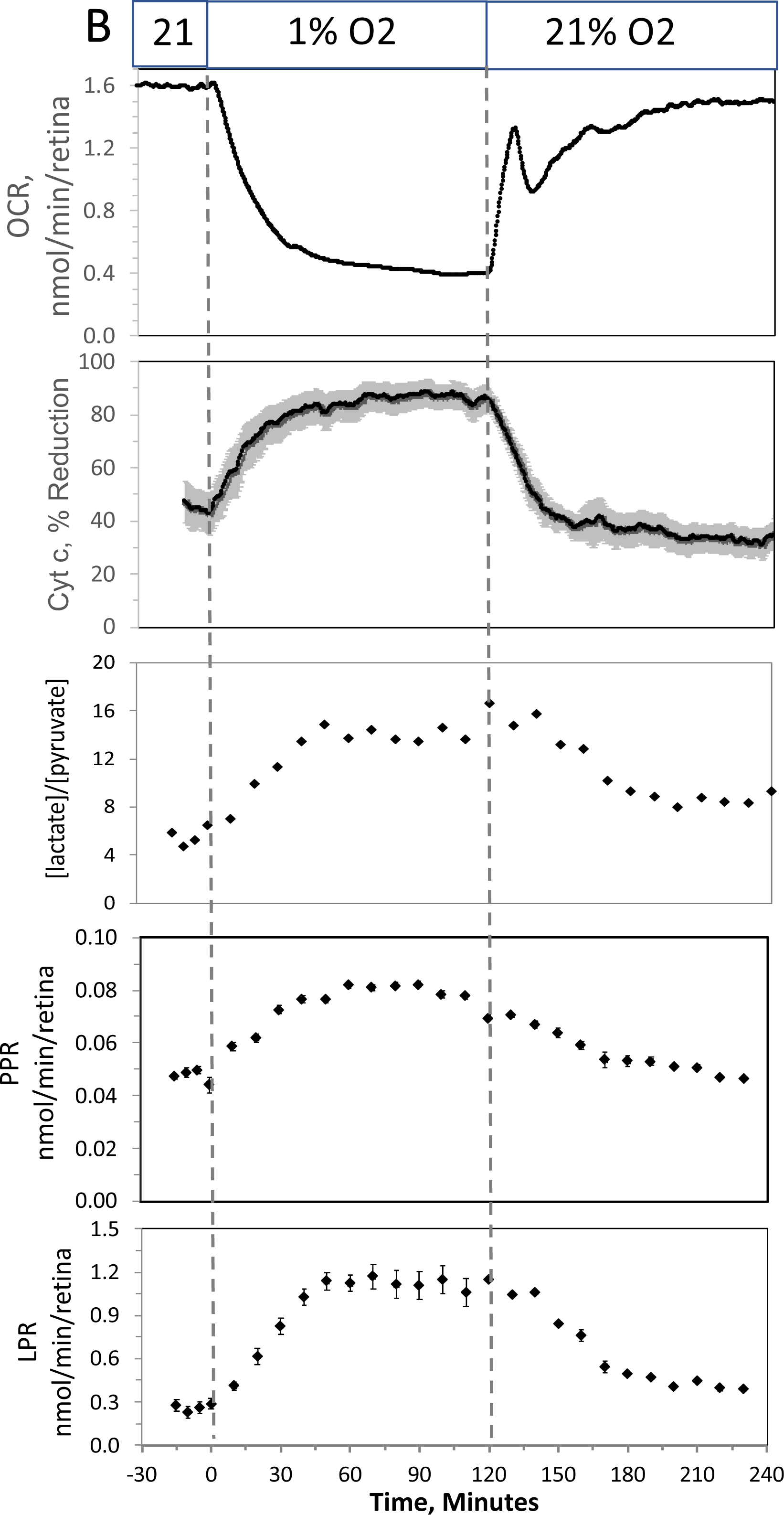
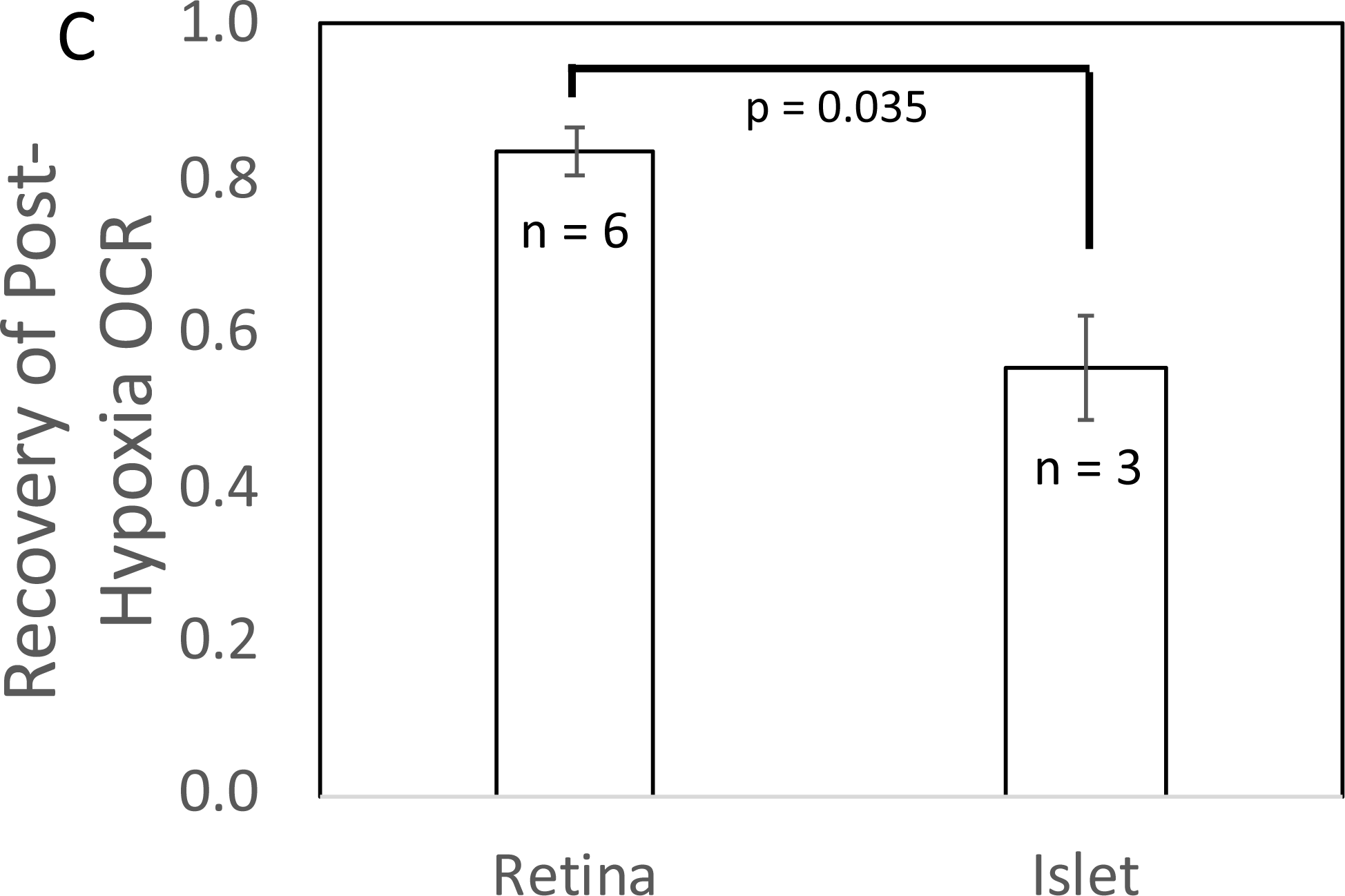
Effect of low O_2_ on transient and steady state ETC, and lactate/pyruvate in isolated retina. **A.** Four retinas (16 pieces) per chamber were loaded into the flow chamber, where each group of 4 retinal pieces were separated by a 3-uL layer of Cytodex beads. The tissue was sandwiched on the top and bottom by 50 uL of Cytopore beads and both layers were held in place with a porous frit (Interstate Specialty Products, Suton, MA, Cat no. POR 4894, cut to 4.2 mm diameter and 0.25 in long). Ninety minutes after loading the retina into the system (flow rate = 130 uL/min), the O_2_ tank was switched one containing 1% O_2_ for 2 hours, and subsequently returned to 21%. **A.** The protocol generated inflow and outflow O_2_ profiles such as shown. Following the completion of the protocol, 12 υg/ml antimycin A (aA) was added for 20 minutes, and then 3 mM KCN, and the hypoxia protocol was repeated while retinal respiration was suppressed in order to characterize delay and dispersion due to the separation in space of inflow and outflow sensors. **B.** Measurements of OCR (representative data from an n of 6 (average recovery = 0.83 +/− 0.03), reduced cytochrome c (n = 6), lactate and pyruvate production rates (n = 2) and [lactate]/[pyruvate] are shown. **C.** Recovery of OCR after hypoxia relative to OCR pre-hypoxia values in retina and islets (t test result: p = 0.035, for retina, n = 6, and for islets n = 3).

In response to hypoxia, lactate and pyruvate production rates by retina increased, consistent with operation of the Pasteur effect. The ratio of lactate/pyruvate also increased during low O_2_ conditions (Fig. 4B), reflecting the decreased uptake of pyruvate into the mitochondria, and the increase in the cytosolic redox state (NADH/NAD) that occurs during low O_2_ ^42^. These results provide support for the utility of measuring extracellular lactate and pyruvate for real time responses to events affecting metabolism

### Complex time-dependent and concentration-dependent effects of H_2_S on ISR by islets resolved by fluidics analysis

Past studies on the effect of H_2_S on ISR were consistent in their findings that H_2_S was inhibitory ^26–30, 43–45^. However, as H_2_S has both stimulatory and inhibitory effects on the ETC ^23, 24^, we predicted that precise titration of the exposure of islets to H_2_S would reveal stimulatory effects of H_2_S on ISR. To simplify the analysis and interpretation, we ramped up the H_2_S concentration in the gas equilibration system (while the inflow and outflow gas ports were clamped) to accumulate H_2_S until the desired concentration was reached, and then clamped the permeation tube inlet port to maintain that H_2_S concentration for the indicated times. When H_2_S was increased until a steady state of 0.44 μM was reached, ISR from pancreatic islets increased by 35% relative to ISR at 20 mM glucose (Fig. 5A). The increased ISR was sustained for 3 hours. The effect of H_2_S was reversible. After purging it from the system, ISR rapidly returned to levels that occurred prior to H_2_S exposure. In the presence of 3 mM glucose, H_2_S had no effect on ISR (data not shown), supporting the idea that this reflects a physiologic response of ISR to H_2_S. To demonstrate the ability of the flow system to more fully characterize the time- and concentration-dependency of ISR on H_2_S, we measured ISR at steady state concentrations of H_2_S from 0.15 to 1.42 μM. Notably, between 0.33 and 1.42 μM H_2_S (Fig. 5B), the initial period of stimulation of ISR (peaking between 1 and 1.5 hours after the start of the ramp of H_2_S) was insensitive to the concentration of H_2_S. In contrast, the effect of higher levels of H_2_S inhibited the ISR rate only between 1.5 and 4 hours following the start of the H_2_S exposure. The initial upslope of ISR occurred with a delay of about 30 minutes, and both the stimulation and inhibition of ISR were rapidly reversible following the washout of H_2_S (Fig. 5B). The ability of the system to resolve the time lag for H_2_S to activate ISR is limited by the rate of the increase of H_2_S in the gas equilibration system accomplished by the permeation tube. In order to increase the temporal resolution, the permeation tube leak rate can be increased, or the volume of the gas equilibration system must be decreased. Additional concentrations of H_2_S were tested and the observed peak of ISR, and the steady state level between 3 and 4 hours were plotted as a function of the concentration of H_2_S (Fig 5C), clearly showing the ability of the system to resolve both the time courses and concentration-dependency of the effects of a relatively small range of [H_2_S]. To test the assumption made in many studies that due to rapid equilibrium between gas donor molecules and their corresponding gas are able to emulate direct exposure to the dissolved gas ^46^, we also analyzed the effect of NaHS on ISR by perifused islets. In contrast to dissolved gaseous H_2_S, low levels of NaHS had no effect, and higher concentrations (>10 μM) only inhibited ISR (Appendix Fig. 1A and 1B) - consistent with findings of all previous studies that used NaHS as an H_2_S surrogate ^26–30, 43–45^. Although it is not clear why there is a difference between effects of H_2_S and NaHS, it may be that the H_2_S generated by NaHS may diffuse out of solution and into the gas phase when the media traverses the gas equilibration system.

**Fig. 5.**
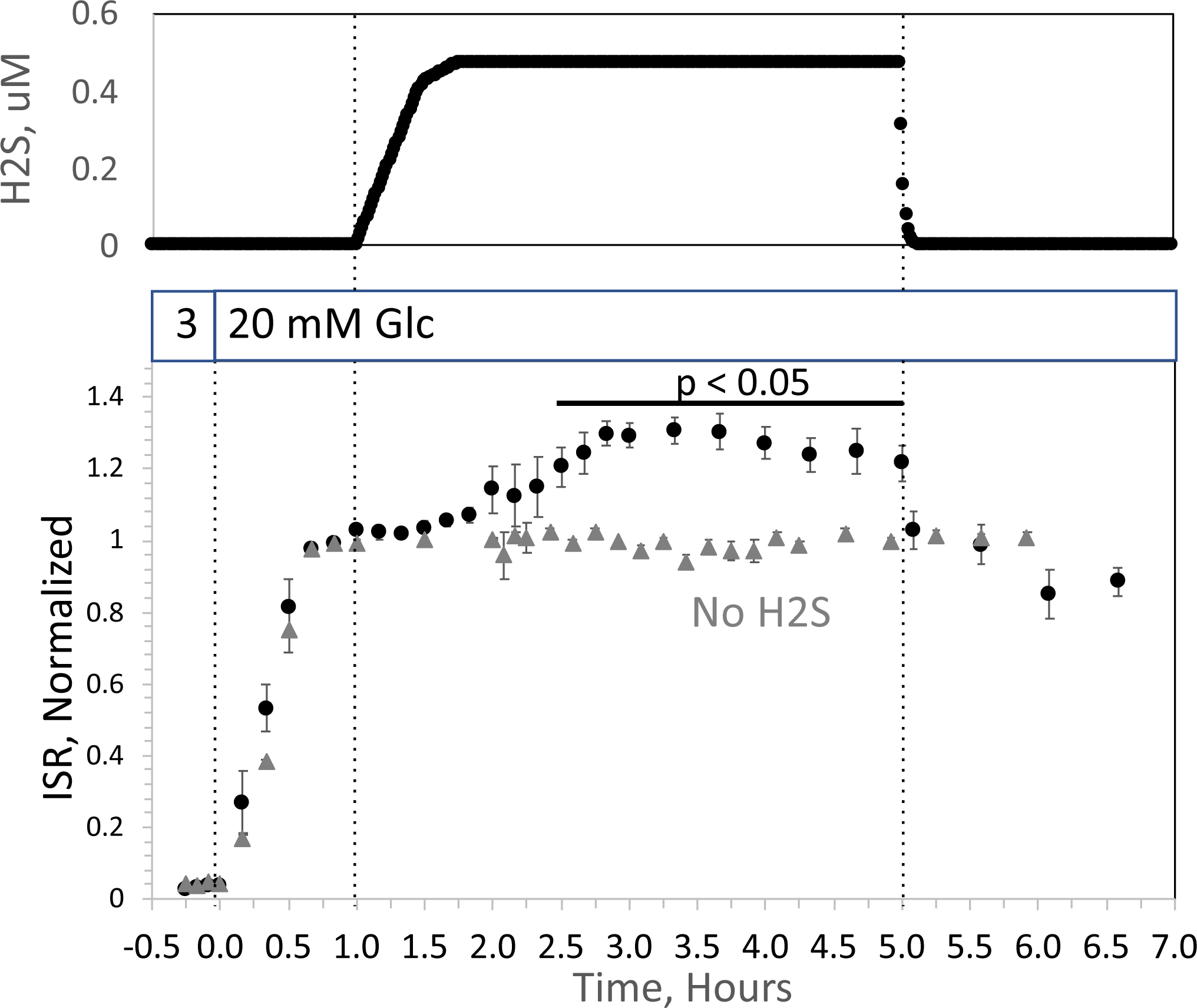
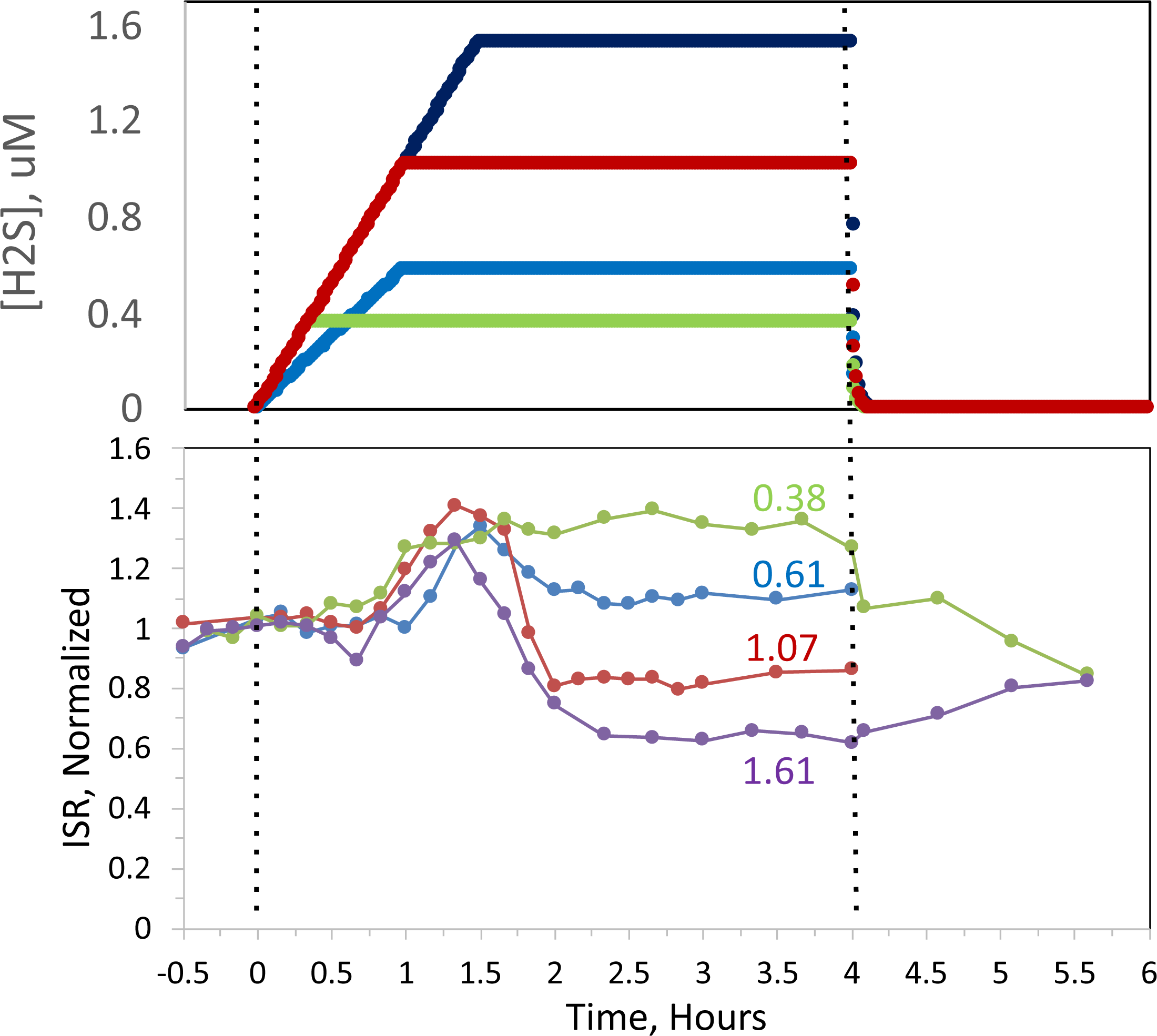
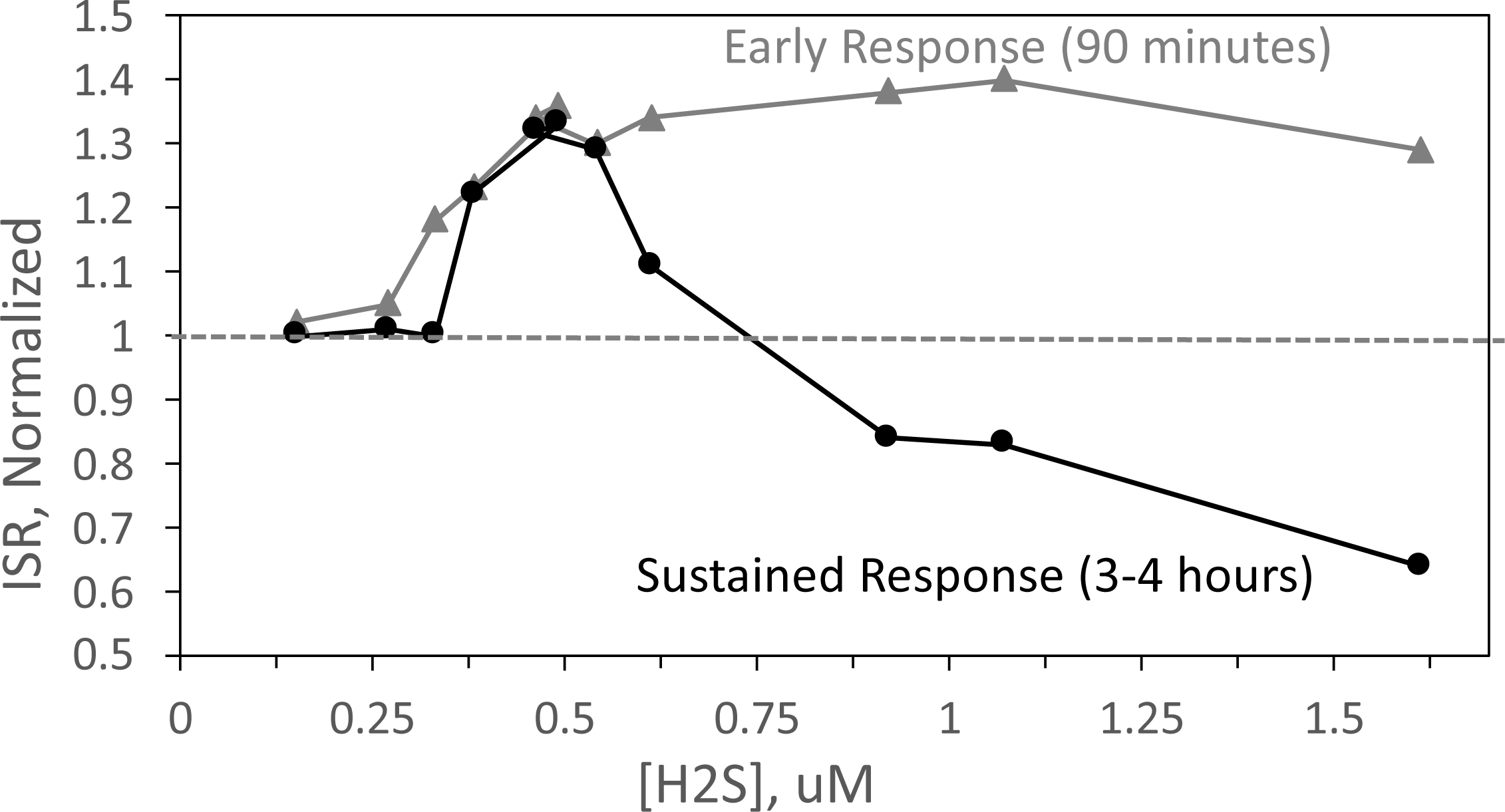
Effect of H_2_S on ISR by islets. **A.** Rat islets (50/channel) were perifused (flow rate = 200 uL/min), and ISR was measured in response to glucose and exposure to dissolved H_2_S in the concentrations shown (data is average +/− SE, n =3 (H_2_S), n = 2 (no H_2_S), p < 0.05 as indicated). **B.** ISR was measured at the indicated concentrations of dissolved H_2_S. Each curve is a single experiment. **C.** Data from perifusions as shown in B were plotted as a function of the ISR at the peak between 1 and 1.5 hours, and the average ISR between 3 and 4 hours.

### Measurement of OCR, cytochrome c, lactate, and pyruvate by perifused liver slices

We also explored the ability of our flow system to measure effects of H_2_S on liver by measuring OCR, cytochromes and lactate/pyruvate release by liver slices in the absence and presence of a mitochondrial fuel (succinate). The responses were complex, changed directions in time- and concentration-dependent fashion, and will ultimately require more experiments to interpret the data mechanistically. Therefore, this data was placed in the appendix (Appendix Fig. 2). Nonetheless the waveforms of the responses were clearly resolved, confirmed the capabilities of the multi-parametric detection system, and so were included in this report. The salient features of the data can be summarized by: 1. In the absence of the mitochondrial fuel succinate, H_2_S changed OCR and reduced cytochromes in proportion to each other, consistent with donation of electrons from H_2_S to cytochrome c ^24^ (Appendix Fig. 2A); 2. In the presence of a TCA cycle intermediate (succinate), H_2_S (2-3 μM) increased the reductive state of cytochrome c oxidase while decreasing OCR. This is consistent with inhibition of cytochrome c oxidase ^23^ (Appendix Fig. 2B), which notably occurred at concentration many times lower than typical estimates of plasma concentration which range from 30-300 μM ^47^. At low levels, H_2_S caused irreversible inhibition of ETC activity upstream of cytochrome c but did not inhibit flow of electrons from succinate. Thus, both reported mechanisms of action of H_2_S on the ETC were resolved by the system, as well as uncovering additional effects of H_2_S that had not previously been reported.

## DISCUSSION

### General features of the flow system

Flow systems have important advantages over static systems for assessment of cell function. Viability and functions of cells and tissues are better and closer to physiological, culture media composition can be changed, and real time production or uptake rates can be quantified from differences between inflow and outflow. Microfluidics devices can maintain tissues in ways that preserve their 3-dimensional structure and preserve native cell to cell interaction^48, 49^. However, these devices will have maximal impact when combined with real time assessment of the tissue as well as the ability to control aqueous and gaseous composition of the media bathing the tissue models. This report focuses on technical modifications to a previously developed flow culture/assessment system ^50^ that enables real time measurements of responses of tissues to physiologically important dissolved gases. We achieved this by incorporating a unique gas equilibration system that controls abundant (blood) gases including O_2_, CO_2_ and N_2_, and by using permeation tubes to introduce and control trace gases such as H_2_S, NO and CO. In this report we demonstrated the utility of this system using both an abundant (O_2_) and a trace gas (H_2_S).

### Control and effects of dissolved O_2_: Recovery of metabolic state following hypoxia and transient response in OCR following reoxygenation

The ability to control dissolved O_2_ makes our system highly suitable for investigating ischemia-reperfusion injury, generating two informative endpoints from a protocol that measures effects of a short period of low O_2_ availability followed by return to normal levels. We used the recovery of OCR and cytochrome c reduction to report tissue sensitivity to hypoxia; and we identified a transient spike in OCR that occurs upon reintroduction of normal O_2_ levels.

The first endpoint characterizes the capacity of a tissue to survive after exposure to selected time periods of low O_2_. In the illustration carried out in this study, retina recovered to 83% after hypoxia, whereas islets recovered to only 55% of pre-hypoxic levels of OCR, corresponding to the two tissue’s known sensitivity to oxidative stress. The second endpoint, the burst of OCR occurring when O_2_ floods back into the cell, is one that has been hypothesized, but has not previously been measured due to the difficulty of measuring OCR in the face of changing O_2_ levels. Our method has enabled the measurement of transient responses to re-oxygenation and revealed a two-phase waveform in OCR in both islets and retina. The ability to resolve this waveform was dependent on rigorous convolution analysis to remove the delay and dispersion of the O_2_ signal due to the flow system, combined with ultra-stable and ultra-sensitive O_2_ sensors. It has been long recognized that ROS is generated rapidly by cells when O_2_ becomes plentiful after undergoing hypoxic conditions ^2, 3^. The transient spike of OCR is consistent with a scenario where O_2_ is the source of oxygen atoms for the ROS. The detailed relationship between OCR and generation of ROS will be the topic of future applications with this system. For instance, the method can be used to test whether slowed re-introduction of O_2_ or candidate therapeutics including H_2_S, prevent or reduce the transient spike in OCR while at the same time preventing the decreased recovery following the hypoxic period (as has been hypothesized ^51–54^). The use of this method is timely given the need for optimization of ventilator settings for recovery of hypoxic bouts in COVID-19 patients ^55^. Other applications could include: testing whether there are differences in transient OCR response for different tissues or metabolic states; determining the relation of the spike to the response of ROS and recovery/survival of tissue; and testing drugs designed to prevent both the transient spike and/or the decrease in post-hypoxic OCR. The ability to objectively quantify recovery of OCR positions the system to be used to test tissue sensitivity to a wide range of stresses (including ER and oxidative stress, immunological stress (exposure to cytokines), lipotoxicity as well as hypoxia), and to test drugs and treatments designed to increase or decrease recovery from experimentally-induced stresses.

### Control and effects of a trace gas: stimulation of ISR by H_2_S in islets

Trace gas signaling molecules (CO, NO, H_2_S) are generated in most tissues and have wide-ranging effects on function, metabolism and protection from hypoxia (for reviews ^56–59^). However, due to the difficulty in quantitatively and reproducibly introducing dissolved trace gases into culture media, the majority of studies on these gases have utilized aqueous chemical donors such as NaHS instead of H_2_S. These surrogates provide only imprecise and uncertain concentrations and timing of tissue exposure to dissolved H_2_S. Due to the extremely accurate calibration of the rate of release of gases, permeation tubes can introduce trace gases into the carrier gas (the mixture of CO_2_-O_2_-N_2_) present in a gas equilibration system of our perifusion apparatus at exact times and concentrations. Producers of permeation tubes can provide them with a selection of over 500 gases, including H_2_S, NO, CO, and ammonia. Thus, the method of incorporating permeation tubes into the gas equilibration system is particularly versatile and can be used for a wide range of applications.

To illustrate the unique advantages of being able to expose tissue to precise levels of dissolved H_2_S, we selected islets as a test tissue since the inhibitory effects of H_2_S on glucose-stimulated ISR by islets have been described. However, those reports were based on use of the H_2_S donors NaHS and Na_2_S ^26–30, 43–45^ and the interpretation that H_2_S gas is actually delivered to the tissue and at levels low enough to avoid its toxic effects. Moreover, we took note of other studies that suggested that H_2_S can donate electrons directly to cytochrome c ^24^ and there is evidence that the reduction of cytochrome c may be a key regulatory step in activating ISR ^34, 60^. Our system bore out this prediction revealing stimulatory effects of H_2_S on ISR that had not been apparent when exposing islets to a donor of H_2_S (NaHS). H_2_S at low concentrations (between 0.25 and 0.5 μM) enhances glucose-stimulated ISR, which remained elevated for at least 4 hours. The concentration- and time-dependency were complex however. At higher concentrations of H_2_S the stimulation of ISR for 90 minutes still occurs, but at later times ISR was inversely proportional to the H_2_S ranging from a 35% increase to a 40% decrease. The range of effects occurred over a relatively small range of H_2_S concentrations, highlighting the need for the very precise control of dissolved H_2_S afforded by the use of permeation tubes to investigate this phenomenon. H_2_S had no effect on ISR at 3 mM glucose, supporting a physiologic mechanism mediating H_2_S’s effect that may be integrated with glucose sensing and secretory response to fuels by the islet ^61, 62^. H_2_S is generated in islets by the action of 3 intracellular enzymes ^63^ but is also a component of blood albeit at levels that are not well established ^64–66^. It is notable that the range of concentrations that induced changes in ISR, and above which caused inhibition of OCR in liver are many times lower than typical estimates of plasma concentration which range from 30-300 μM ^47^. Although it is possible that in vivo, tissue is much less sensitive to H_2_S than in vitro, it seems more likely that assays that measure plasma H_2_S result in a significant overestimate coming from related sulfur-containing compounds. The ability to detect differences between responses to H_2_S and donor molecules, will be useful to the increasing numbers of investigators developing H_2_S donor molecules as pharmaceutics ^67, 68^. The increase in ISR in response to H_2_S has physiological, methodological and clinical implications and the lack of similar stimulatory effects by a donor molecule has broad implications in a field where studies of NO, H_2_S and CO are mostly based on the use of donor molecules.

### Lactate and pyruvate: relation to cytosolic events

The assayed values of lactate and pyruvate reflect a number of important specific and global parameters. The rate of release of lactate and pyruvate is an integration of the rate of glycolysis less the amount of pyruvate flux into the mitochondria and traversing gluconeogenesis. Thus, both compounds generally increase in response to glycolytic fuels. Importantly for the study of hypoxia, both metabolites rise in cells when O_2_ is decreased (the Pasteur effect). In addition, the ratio of cytosolic lactate/pyruvate mirrors the cytosolic NADH/NAD ratio due to the equilibrium status of the LDH reaction ^69^. One could envision that freeze clamping cells and measuring intracellular lactate and pyruvate to directly compare them to the extracellular values could be a way to validate the use of extracellular data. However, in practice, the measurement of intracellular compounds is difficult and also limited by the kinetic resolution of freeze clamping. Instead, we evaluated the responses of the extracellular levels of lactate to a blocker of LDH (oxamate), of pyruvate to a blocker of mitochondrial transport (zaprinast), and both compounds in response to hypoxia. The rapid changes in extracellular lactate and pyruvate supports the rapid redistribution between intra- and extra-cellular compartments, and that real time measurement of extracellular lactate and pyruvate reflect intracellular events governing intracellular lactate and pyruvate. Retina responded to hypoxia with a classical Pasteur effect: low O_2_ increased lactate, pyruvate and lactate/pyruvate ratio. We envision that the measurement would be especially informative when examining the shift from oxidative to glycolytic metabolism such as seen in tumorigenesis ^70^ or stem cell differentiation^71^.

### Incorporation of O_2_-control into a real time fluorescent imaging system

Real time fluorescent imaging is a powerful modality that is commonly used to quantify a wide variety of intracellular compounds and factors while perfusing the optical chamber housing the cells or tissue. Molecular Probes provides intracellular dyes for over 100 separate compounds, so this method is versatile and wide-ranging. Thus, incorporating the gas equilibration system to a flow system providing buffer to a chamber that images of single islets. We observed and quantified clear increases in intracellular Ca^2+^ in response to hypoxia as metabolic rate decreased. Unexpectedly, the loss of energy and ISR following islet exposure to hypoxia were not accompanied by a loss of glucose-stimulated Ca^2+^, suggesting the mechanism mediating loss of ISR is independent of Ca^2+^ signaling. These data are consistent with previous findings that loss of secretory function is more closely associated with bioenergetics than Ca^2+^ ^34, 72^. When comparing results of various assays and modes of analysis, the ability to measure multiple endpoints under matched conditions and the same flow system is optimal for systematic study of tissue function.

### Summary of uses for the flow culture system

The ability to precisely control the levels and timing of exposure to both abundant and trace gases while measuring multiple parameters in real time on a wide range of tissue and cell models make this system uniquely powerful. The novel resolution of OCR transients attests to the high kinetic resolution of the system. Moreover, stimulatory effects of H_2_S on ISR not seen in response to a donor molecule attests to the ability of the technology to reveal behavior that provides new insight. Given the wide use of donor molecules in studying gasotransmitters this has broad and significant implications. In addition to enabling users to evaluate direct effects of gases, the method is also suitable for testing conditions or agents that diminish loss of OCR in response to hypoxia- or other stress-induced effects. The use of methods to study the effects of dissolved gas on tissue will impact many areas of fundamental research as well as research of diseases including but not limited to diabetic wound healing, stroke (ischemia/reperfusion injury), COVID-19 and cancer.

## METHODS

### Chemicals

Krebs-Ringer Bicarbonate buffer was used for all perifusions prepared as described previously^12^. Antimycin A, glucose, oxamate, KCN and zaprinast were purchased from Sigma-Aldrich. Gases of varying O_2_ levels/5% CO_2_ and balance N_2_, were purchased from Praxair Distribution Inc (Danbury CT). Cytodex and Cytopore beads were purchased from GE healthcare and biosciences (cat no. 17-0448-01 and 17-0911-01, respectively).

### Culture of INS-1 832/13 cells

INS-1 832/13 cells were kindly provided by Dr. Christopher Newgard and were cultured as previously described ^73^. The day before experiments, cells were harvested, and cultured with Cytodex beads (2.5 mg/million cells) for 15 min in RPMI Media 1640 (Gibco, Grand Island, NY) supplemented with 10% heat-inactivated fetal bovine serum (Atlanta Biologicals, Lawrenceville, GA), 2 mM L-Glutamine, 1 mM Pyruvate, 50 μM Beta-mercaptoethanol, 20 mM HEPES and 1% Pen/Strep. They were then washed and cultured overnight in a standard CO_2_ incubator at 37 degrees C.

### Tissue harvesting and processing

All procedures were approved by the University of Washington Institutional Animal Care and Use Committee.

#### Rat islet isolation and culture

Islets were harvested from male Sprague-Dawley rats (approximately 250 g; Envigo/Harlan, Indianapolis, IN) anesthetized by intraperitoneal injection of sodium pentobarbital (150 mg/kg rat)) and purified as described ^50, 74^. Subsequently, islets were cultured for 18 hours in RPMI Media 1640 supplemented with 10% heat-inactivated fetal bovine serum (Invitrogen) at 37°C prior to the experiments.

#### Retina isolation

Retinas were harvested from C57BL/6J mice (euthanized by cervical dislocation) ten minutes prior to loading and were dissected into ¼ths using micro scissors as previously described ^16^.

### Flow Culture System to maintain tissue with precise control of dissolved gases

A flow culture system^12^ was modified to continuously perifuse tissue with buffer equilibrated with the desired composition of dissolved gas using a gas-equilibration system (Fig. 6A and B). Multiple modes of assessment were integrated into the flow culture system and are described below including chemical sensors for O_2_, spectroscopic analysis of the tissue for measurement of reduced cytochrome c and cytochrome c oxidase, and collection of outflow fractions for subsequent assay of lactate and pyruvate, or insulin. Model numbers and manufacturers are listed in the legend for Fig. 6. Prior to entering the perifusion chamber, perifusate is pumped from the media reservoirs by an 8-channel peristaltic pump into the thin-walled silastic tubing of the gas equilibration system that facilitated equilibration between the buffer and gas in the glass housing. Selection of media reservoir with desired media composition determined by use of a 6-port valve. To achieve the desired gas composition of O_2_, CO_2_ and N_2_ within the gas equilibration system, tanks of premixed gases supplied gas to the inflow port, typically 5% CO_2_, the desired percentage of O_2_, and balance N_2_.

**Fig. 6.**
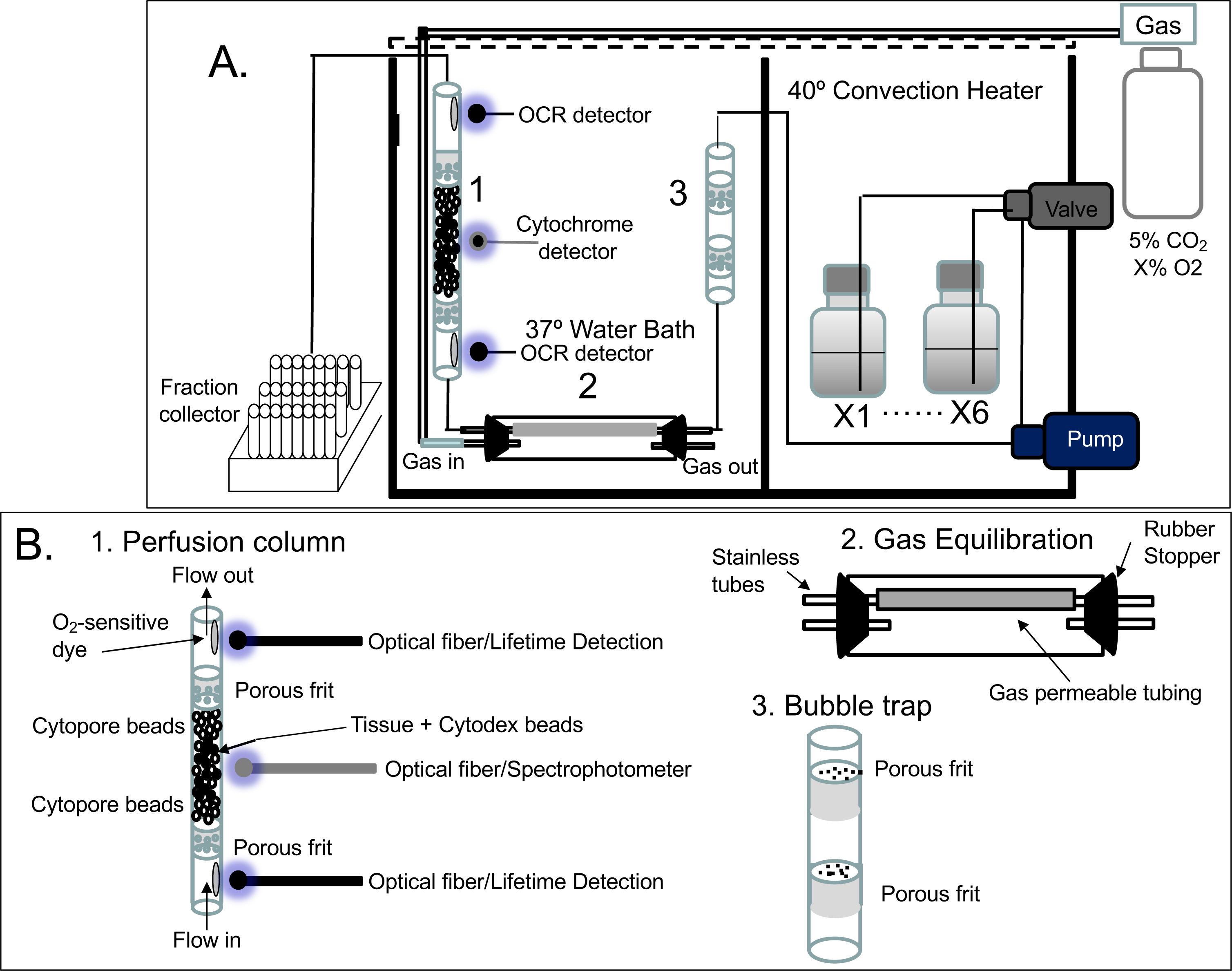
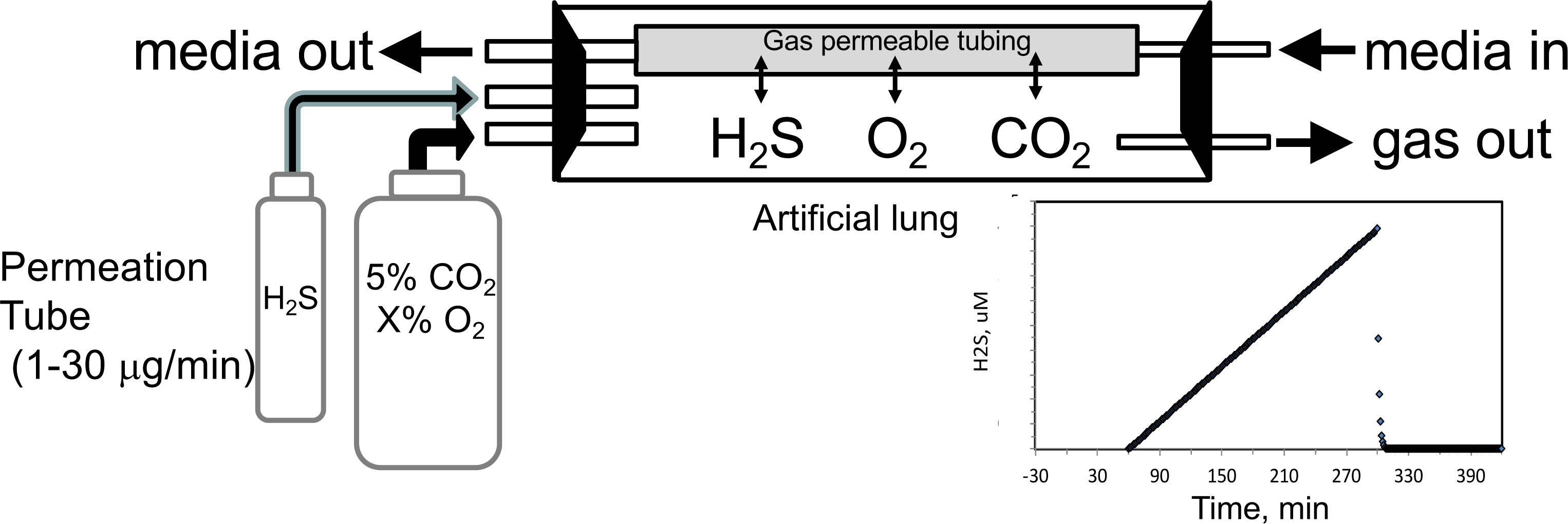
Flow culture system/assessment system for studying effects of dissolved gases on tissue or cells. One channel/perifusion chamber is shown, but the actual system can accommodate up to 8. **A.** The perifusion system consisted of an 8-channel peristaltic pump (MiniPuls 2, Gilson, Middleton, WI) connected to a 6-port valve (Part # V-451, IDEX Health and Science, Oak Harbor, WA) to produce up to six separate solutions; a media/dissolved gas equilibrium system in which media flowed through thin-walled Silastic™ tubing (0.062 in ID x 0.095 in OD; Dow Corning Corp., Midland, MI) for a residence time of 5 minutes (typically 0.2-0.5 m depending on the flow rate) loosely coiled in a glass jar that contained various O_2_, 5% CO_2_/balance N_2_; bubble trap comprised of a Simax Borosilicate glass tube (Mountain Glass, Asheville, NC, 2” long and 4.2 mm ID) filled with glass wool; a Simax Borosilicate glass perifusion chamber (3” long and 4.2 mm ID) immersed in a 37°C water bath; Lifetime detection spectrometers (Tau Theta, Boulder CO), a tungsten-halogen light source/USB2000 spectrophotometer (Ocean Optics OH); and a Foxy 200 fraction collector (Isco, Inc., Lincoln, NE). **B.** A blow up of individual parts shown in A. 1.) the glass perifusion chamber containing culture beads, porous frits (Interstate Specialty Products, Suton, MA, Cat no. POR 4894, cut to 4.2 mm diameter and 0.25 in long) to support the tissue and disperse the flow, and coated with O_2_-sensitive dye on the interior above and below where the tissue resides; 2) gas equilibration chamber, where media flows through gas permeable Silastic™ tubing and equilibrates with the gasses filling the headspace; 3) bubble traps. **C.** Incorporation of a permeation tube (VICI Metronics, Poulsbo, WA) that releases H_2_S at specified rates into the media/dissolved gas equilibration system during which time the ports for the inflow and outflow of carrier gas (O_2_, C O_2_ and N_2_) are closed. The resulting accumulation of H_2_S in the artificial lung yields linearly increasing concentrations of dissolved H_2_S in the form of a ramp function as shown.

To equilibrate the inflow with desired concentration of H_2_S, we have used devices called permeation tubes (VICI Metronics, Poulsbo WA), an industry standard that provides very precise rates of gas release, typically from 1-30 ug/min. The outlet of the permeation tube was connected to the inlet port of the chamber housing the gas equilibration system (Fig. 6C), so that the head space around the perifusate in the gas-permeable tubing accumulated the trace gas. The concentration in the chamber then increased as a ramp function, rising at a rate equal to the leak rate of the permeation tube x time divided by the volume of the housing. The amount of dissolved H_2_S in the buffer was then calculated based on the solubility of H_2_S in buffer based on Henry’s Law:

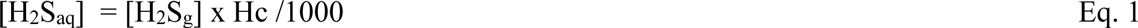

where Henry’s constant Hc is in 0.1 atm/M, [H_2_S_aq_] is in μM, and [H_2_S_g_] is in ng/mL

#### Lifetime detection of dissolved O_2_

O_2_ tension in the inflow and outflow buffer was measured by detecting the phosphorescence lifetime of an O_2_-sensitive dye painted on the inside of the perifusion chamber using a MFPF-100 multifrequency phase fluorometer lifetime measurement system (TauTheta Instruments, Boulder, CO) as previously described^75^. Using tanks of gas containing varying amounts of O_2_ (21, 15, 10, 5, 3, 1 or 0%), data was generated that showed the dependency of the lifetime signal as a function of O_2_ and the rapidity of changes in O_2_ after each change in gas tank. Within 5 minutes, O_2_ achieves 95% of steady state levels (Fig. 7A), where the delay is primarily due to time needed for the gas in the gas equilibration system to turnover as the actual sensor responds in microseconds. The O_2_ dependency of the dye signal conformed to the Stern-Volmer equation

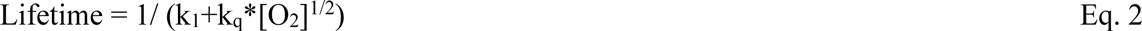

where lifetime is in usec. Equation 2 was used to as a calibration curve to convert the optical signals to O_2_ content (Fig 7B). The use of lifetime detection produces very stable and sensitive data at both normal (Fig. 7C) and low (Fig. 7D) O_2_ levels producing S/N over 20 even when measuring a change of only 1.9 mm Hg.

**Fig. 7.**
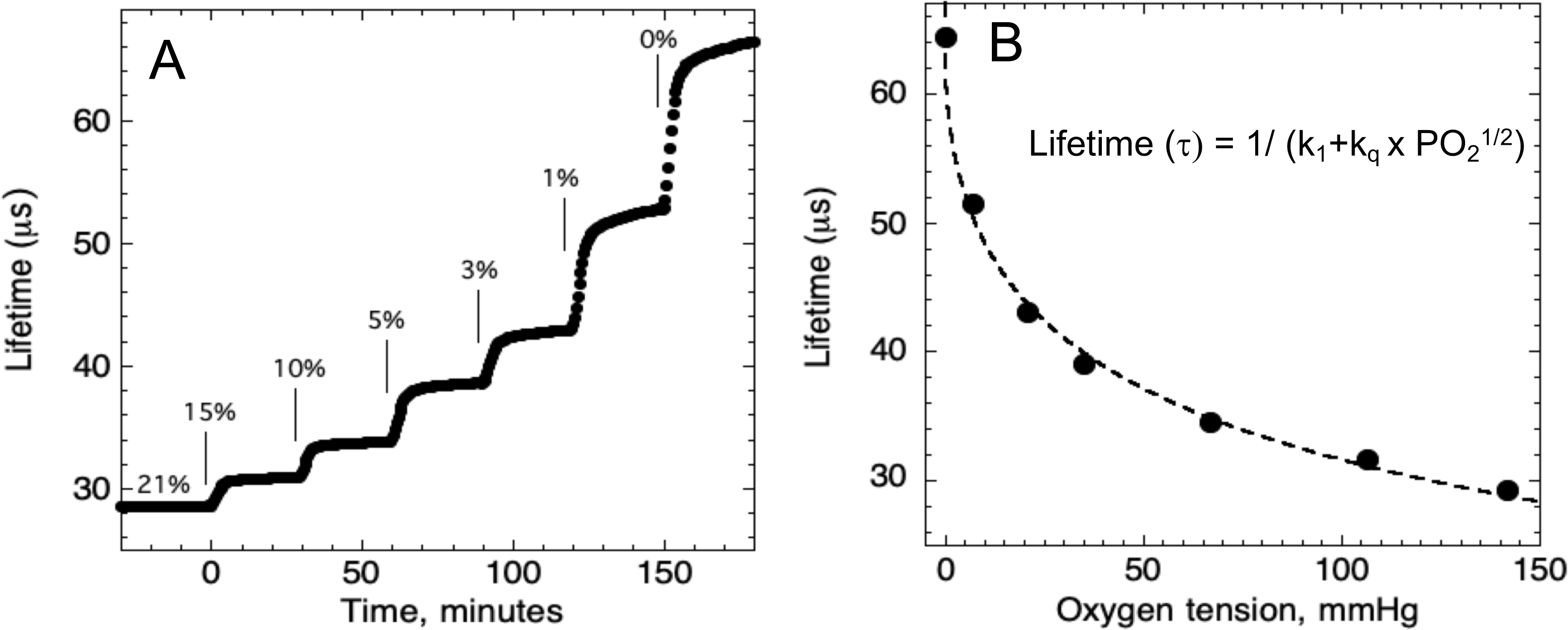
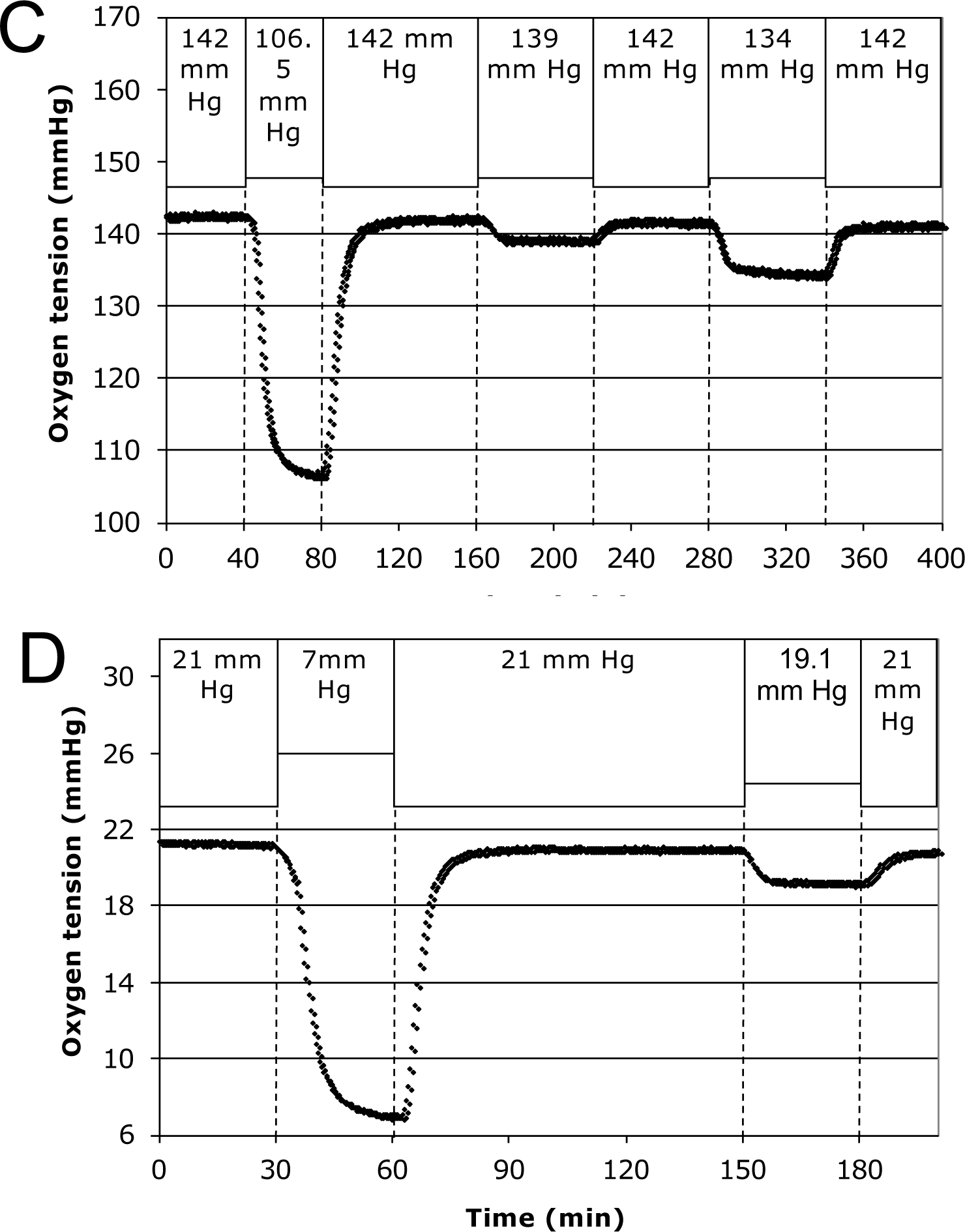
Control and measurement of dissolved O_2_. **A.** At 20-30 minute intervals, the lung was sequentially filled with 21, 15, 10, 5, 3, 1, and 0% O_2_. **B.** The steady state lifetime measurements were non-linear and conformed to a Stern-Volmer equation. Where the pO_2_ is partial pressure of oxygen molecules (oxygen tension), τ0 is lifetime of the dye without quencher such as oxygen molecule, k_1_ is 1/τ0 and k_q_ is a bimolecular quenching constant of the dye by oxygen molecules. The k_1_ and k_q_ were 15519 sec^-1^ and 1612.2 mmHg ^-1^ sec^-1^, respectively. **C.** Test of precision at normal O_2_. O_2_ (142 mm Hg) in the lung was changed first to 106.5mm Hg and back to 142 (levels of change that were typical what was observed in our studies). By mixing 21 and 15% tanks at known flow rates, O_2_ was then decreased by 3 mm Hg and then subsequently by 8 mm Hg. **D.** Test of precision and S/N of low O_2_. O_2_ (21 mm Hg) in the lung was changed first to 7 mm Hg (1% O_2_) and back to 21. By mixing 3 and 1% tanks at known flow rates, O_2_ was decreased by 1.9 mm Hg. S/N was for the 14 mmHg and the 1.9 mm Hg changes was >80 and > 10 respectively.

### Continuous measurement of OCR

#### Measuring the difference between inflow and outflow during invariant inflow O_2_

When inflow O_2_ tension is constant, OCR by the tissue equals the difference between the content of O_2_ flowing into the perifusion chamber minus that flowing out time the flow rate as follows.

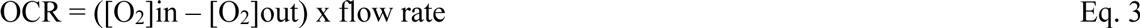

where flow rate is in uL/min and [O_2_] is in nmol/mL. Inflow and outflow O_2_ sensors were positioned on the inside of the perifusion chamber 2 cm upstream and 2 cm downstream from the tissue, respectively. Perifusate flow rates were set to result in a difference between inflow and outflow O_2_ of between 5 and 25% of the baseline O_2_ signal, so it was large enough to be accurately measured, but small enough to avoid exposure to unintended hypoxic conditions.

#### Measuring OCR during changes in inflow O_2_: convolution analysis to remove system effects

Measuring the temporal changes in OCR by tissue in the face of changing inflow concentrations of O_2_ requires a correction for the difference in inflow and outflow O_2_ levels due to the delay and dispersion generated between the inflow and outflow sensors. To calculate OCR from equation 3 in the face of changing inflow O_2_, the inflow O_2_ content must be converted to what it would be if the sensor was located at the outflow sensor location. This was done with classical convolution methods ^76^ with mild regularization^77^ to create a mathematical function representing the delay and dispersion of the inflow signal by the flow through the perifusion chamber from the inflow to the outflow sensor described numerically by equation 4.

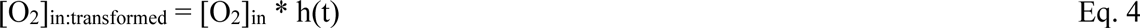

where [O_2_]_in:transformed_ is the inflow concentration at the outflow sensor, and h(t) was the system transfer function. In the absence of tissue in the flow system, O_2_ was decreased to hypoxic levels in the same protocol as was done in the presence of tissue, while measuring [O_2_] in the inflow and outflow (Fig. 8). The transfer function h(t) was then generated for each experimental condition by solving equation 4 by deconvolution using MatLab. For each perifusion analysis, the measured [O_2_]_in_ was converted to [O_2_]_in:transformed_ by convolution with the transfer function (also using MatLab) and OCR was calculated from

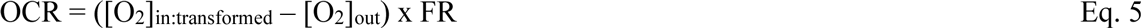

**Fig. 8.**
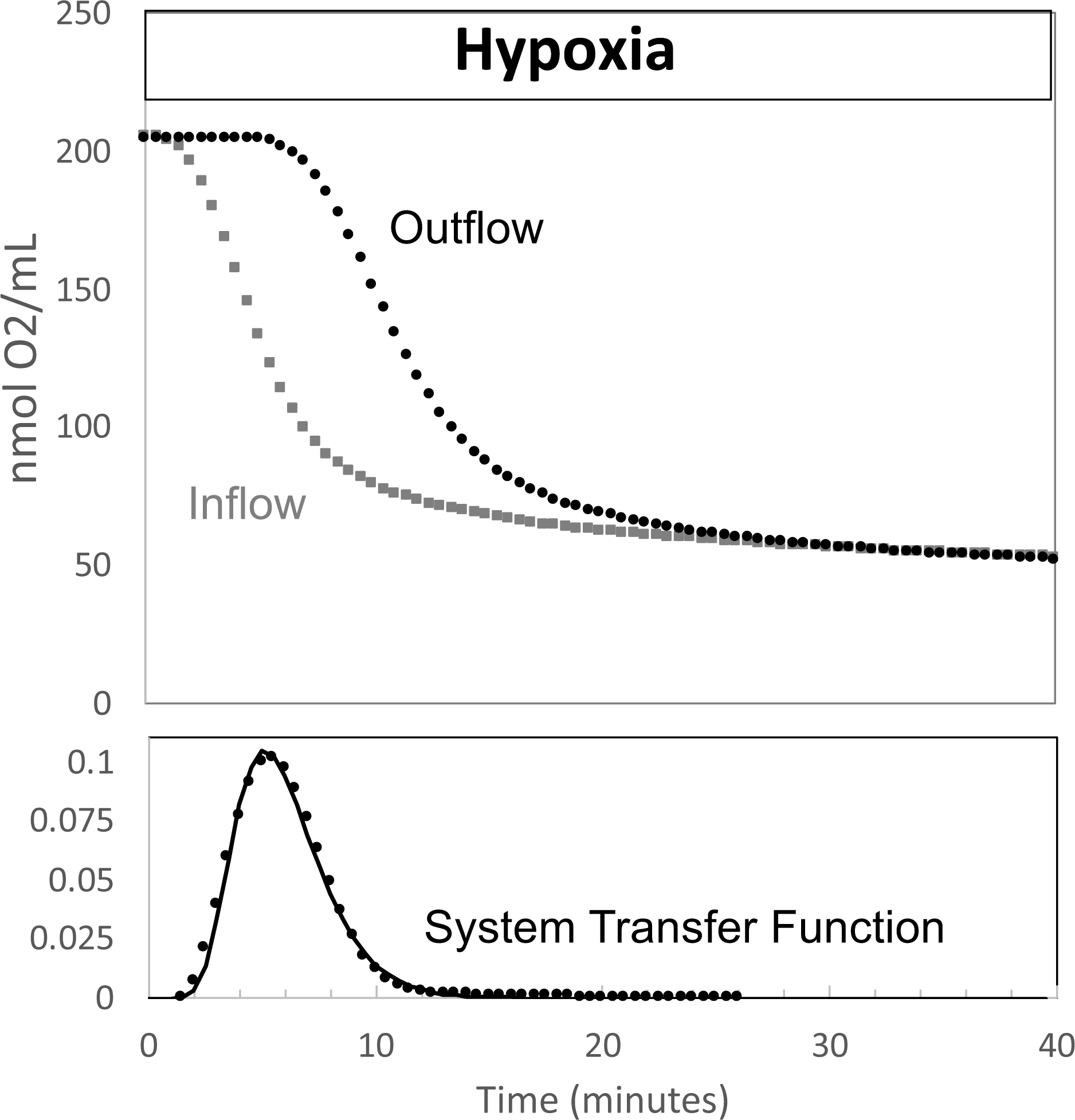
Determination of transfer function from inflow and outflow for convolution analysis. Measurement of inflow and outflow O_2_ tensions in response to a change from 21 to 1% with no live tissue in the system. Deconvolution was carried out to generate the transfer function of the system shown on the bottom graph.

For the protocols in this experimental set-up, only 35 minutes of the transfer function was needed to accurately transform the inflow [O_2_].

### Measurement of cytochrome c and cytochrome c oxidase reduction

The reductive states of cytochrome c and cytochrome c oxidase were measured by light transmission at 550 and 605 nm respectively through the column of islets or tissue as previously described ^60, 78^. Due to the low signal to noise and baseline shift during the experiments, direct measurement of absorption was not stable. To better resolve changes in the reduced state of cytochromes the second derivative of the absorbance spectra with respect to wavelength was calculated^79^. Like absorption, this parameter reflects the number of electrons bound to the cytochrome as well as the amount of protein. However, the second derivative is unaffected by shifts in baseline allowing resolution of real time changes in absorbance. At the conclusion of each experiment, calibration spectra for fully oxidized and reduced cytochromes were acquired in the presence of blockers of the ETC - namely 12 ug/ml antimycin to stop the flow of electrons to cytochromes, followed by 3 mM KCN to facilitate the maximal accumulation of electrons bound to cytochromes.

#### Spectral data processing

The second derivative of absorbance with respect to wavelength^79^ at 550 and 605 nm was calculated as

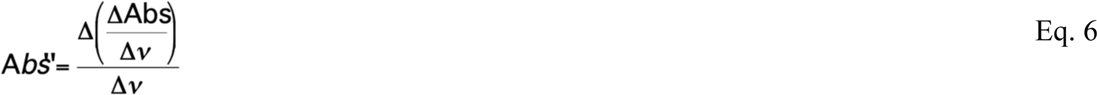

where Abs = log (intensity - intensity_bkg_) / (intensity_ref_ - intensity_bkg_), *v* = wavelength in nanometers, and Δ = change in the variable over the integration interval. Background intensity (intensity_bkg_) was determined with the light source off, and the reference intensity (intensity_ref_) was that obtained when cytochromes were fully oxidized by antimycin A. Percent reduction of cytochromes were calculated following Kashiwagura et al.^80^ as

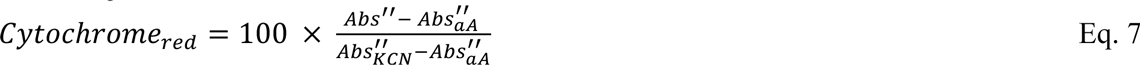

where Abs’’ and Abs’’_KCN_ are values at 550 or 605 nm, and Abs’’_KCN_ and Abs’’_aA_ is obtained in the presence of KCN and antimycin A corresponding to when cytochrome c and cytochrome c oxidase are fully reduced or oxidized.

### Assays for lactate, pyruvate and insulin

Fractions collected during experiments were subsequently assayed for lactate, pyruvate or insulin. Insulin was measured by RIA, and lactate and pyruvate were measured using colorimetric assays using kits per manufacturer’s instructions (insulin, Cat no. RI-13K, Millipore Sigma, Burlington, MA; lactate, Cat no. A22189, Invitrogen, Carlsbad, CA; pyruvate, Cat no. MAK332, Sigma Aldrich). Amounts of lactate, pyruvate and insulin in inflow samples were insignificant, so rates of production were calculated as the concentration in the outflow times the flow rate and normalized by the amount of tissue.

### Imaging and quantification of cytosolic Ca^2+^

Cytosolic Ca^2+^ was measured by fluorescence imaging of islets after loading them with Fura-2 AM (Invitrogen) as previously described ^81^. The perifusion system described above was used to supply buffer with the specified gas composition to a temperature-controlled, 250-μl perifusion dish (Bioptechs, Butler, PA) that was mounted on to the stage of a Nikon Eclipse TE-200 inverted microscope. Results are displayed as the ratio of the fluorescent intensities during excitation at two wavelengths (F340/F380).

### Statistical Analysis

When the message to be conveyed by the graph was an illustration of the high resolution and low noise of the data that was generated by the method, then single experiments were shown as indicated, for instance for OCR in response to hypoxia. In most instances, to demonstrate the reproducibility of the data, technical replicates were conducted and the data averaged - i.e. multiple perifusion channels were run in parallel with pooled tissue or cells batches from multiple animals or flasks of cells. When the goal was to test and show a biological effect, multiple runs were done on different days (for instance comparison of retina and islet recovery of OCR following hypoxia) and statistical significance was determined using Student’s t tests carried out with Microsoft Excel (Redmond, WA). With either technical or biological replicates, error bars on time courses were calculated as the average +/− the SE (calculated as SD/n^1/2^). Raw data for all experiments is compiled into an Excel spreadsheet and saved as a source file.

## ABBRIEVIATIONS

CO_2_: carbon dioxide
CO: carbon monoxide
ETC: electron transport chain
H_2_S: hydrogen sulfide
ISR: insulin secretion rate
KCN: potassium cyanide
LDH: lactate dehydrogenase
MPC: mitochondrial pyruvate carrier
NaHS: sodium hydrosulfide
NO: nitric oxide
O_2_: oxygen
OCR: oxygen consumption rate

## ACKNOWLEDGEMENTS

This research was funded by grants from the National Institutes of Health (DK17047) and the National Science Foundation (STTR Phase 2, 1853066). Special thanks to VICI Metronics for providing custom-made permeation tubes.

## AUTHOR CONTRIBUTIONS

IRS devised the methods and designed the experiments with contributions for retinal experiments from JBH. JK contributed to the incorporation of permeation tubes into the gas equilibrium system. VK, BMR and SRJ performed the experiments; KPB developed the data processing algorithms for calculation of OCR; IRS conceived the study and wrote the manuscript with helpful feedback from JBH. All authors read and commented on the final manuscript.

## APPENDIX

### APPENDIX METHODS

#### Chemicals

Antimycin A, glucose, potassium cyanide (KCN) and succinate were purchased from Sigma-Aldrich.

#### Preparation of Liver slices

Liver pieces from rats were prepared as previously described^1^. Ten pieces (about 0.25 × 1 mm (mass= 2.5–3.5 mg per piece)) were loaded into each tissue perifusion chamber without succinate, or four pieces for chambers with succinate for each analysis. After the end of each experiment, liver samples were weighed, and OCR measurements were normalized to this mass. Liver pieces had no Cytodex bead layering, but otherwise had the same loading procedure as described for retina chambers, the same supplemented KRB was continuously flowed at a rate of about 95 uL/min after initially loading the tissue at 30 uL/min. Procedures were approved by the University of Washington Institutional Animal Care and Use Committee.

### APPENDIX RESULTS

#### Effect of NaHS on glucose-stimulated ISR by isolated rat islets

To test the ability of NaHS to emulate the of H_2_S, islets in the flow culture system at incrementally increasing concentrations of NaHS. No effect was seen at concentration below 10 μM (APPENDIX Fig. 1A), a range that H_2_S had both stimulatory and inhibitory effects on ISR. At higher concentrations, NaHS inhibited ISR where the IC50 was about 30 μM and each concentration change reached a new steady state within 10-15 minutes.

#### Effect of H_2_S on liver in the presence and absence of a mitochondrial fuel (succinate)

To test the ability of our system to measure the effects of H_2_S, rat liver pieces were placed into the perfusion chambers and exposed to steadily increasing concentrations of dissolved H_2_S. The first effects observed in response to changes in H_2_S were the increased production rates of lactate and pyruvate (APPENDIX Fig. 2A). About 30 minutes following the start of the ramp increase in H_2_S, OCR, reduced cytochrome c and cytochrome c oxidase all decreased very precipitously for about 20 minutes, followed by a sudden increase until H_2_S was purged from the system (APPENDIX Fig. 2A). During the post-H_2_S phase, all three of these parameters remained low. After a second introduction of H_2_S, OCR, reduced cytochrome c and cytochrome c oxidase all increased about 30 minutes following the start of the ramp. These waveforms suggest multiple points of action by H_2_S and is consistent with a scenario where H_2_S can irreversibly inhibit complex 1 or a step upstream from complex 1 (consistent with decreased cytochromes and OCR), but also can supply electrons directly to cytochrome c when the level of H_2_S reaches higher concentrations.

In the presence of a fuel that enters at complex 2 (succinate), OCR, reduced cytochrome c and reduced cytochrome c oxidase all increased (APPENDIX Fig. 2B). Under these conditions, H_2_S decreased OCR and cytochrome c reduction, but increased cytochrome c oxidase reduction, suggesting that H_2_S was inhibiting both at complex 1 and 4. In contrast to the experiments done in the absence of succinate, post-H_2_S metabolism was only slightly different than pre-H_2_S energy state reflecting the continued supply of electrons to the ETC when complex 1 is still inhibited, while complex 4 inhibition by H_2_S seemed to be reversible. Following washout of both H_2_S and succinate, reducing power in the mitochondria fell to near 0, as would be expected in the absence of complex 1 activity.

### APPENDIX DISCUSSION

#### Contrasting effect of H_2_S and NaHS on ISR in the perifusion system

The assumption made in most studies of H_2_S, is that NaHS or Na_2_S would rapidly equilibrate with the protonated form of the acid. Whether NaHS or H_2_S is added to the solution, the same amount of H_2_S would be present in solution after a short equilibration time. That H_2_S stimulated ISR and NaHS did not is therefore hard to explain. One factor to consider is that as the buffer traversed the lung, a significant proportion of the H_2_S is depleted from the solution as it diffuses into the gas in the head space around the gas-permeable tubing in the gas equilibration system. This would likely be the case in the static conditions that ISR assays are commonly carried out. Nonetheless, one would reason that as NaHS is increased, eventually the dissolved H_2_S would reach an amount that would stimulate ISR. We have no explanation for why this did not occur, but possibly as HS^-^ gets high enough it can inhibit ISR. So, despite the fact that we cannot explain the lack of stimulatory effect of NaHS, it does support that when investigating the physiological effects of H_2_S, dissolved H_2_S should be used and calls into question the use of H_2_S donor molecules.

#### Control and effects of H_2_S on liver

To illustrate its use, we exposed liver to H_2_S, an established cell signal with multiple effects and mechanisms of action. Both reported mechanisms of action of H_2_S in the ETC were observed in this study: inhibition of cytochrome c oxidase ^2^ and direct transfer of electrons to cytochrome c ^3^.

As the concentration of dissolved H_2_S increased, reduced cytochrome c, reduced cytochrome c oxidase and OCR all decreased reflecting indicating inhibition of step(s) upstream of complex 1 (which appeared irreversible), and subsequently all three parameters increased consistent with donation of electrons directly from H_2_S. In the presence of a mitochondrial fuel entering at complex 2 (succinate), H_2_S caused a decrease in reduced cytochrome c and OCR, while increasing reduced cytochrome c oxidase consistent with the simultaneous inhibition of complex 1 and 4. This analysis highlights the benefit of measuring reductive states of cytochromes concomitantly with OCR: interpreting observed changes in OCR in terms of the mechanisms mediating the changes is not possible with OCR measurements alone. The factors mediating the changes in OCR is distinguished by concomitant measurement of both electron pool sizes in the ETC (cytochromes) and flux of electrons (OCR). Overall, the method was able to characterize complex and multiple effects of H_2_S on the ETC.

---

**APPENDIX Fig. 1.**
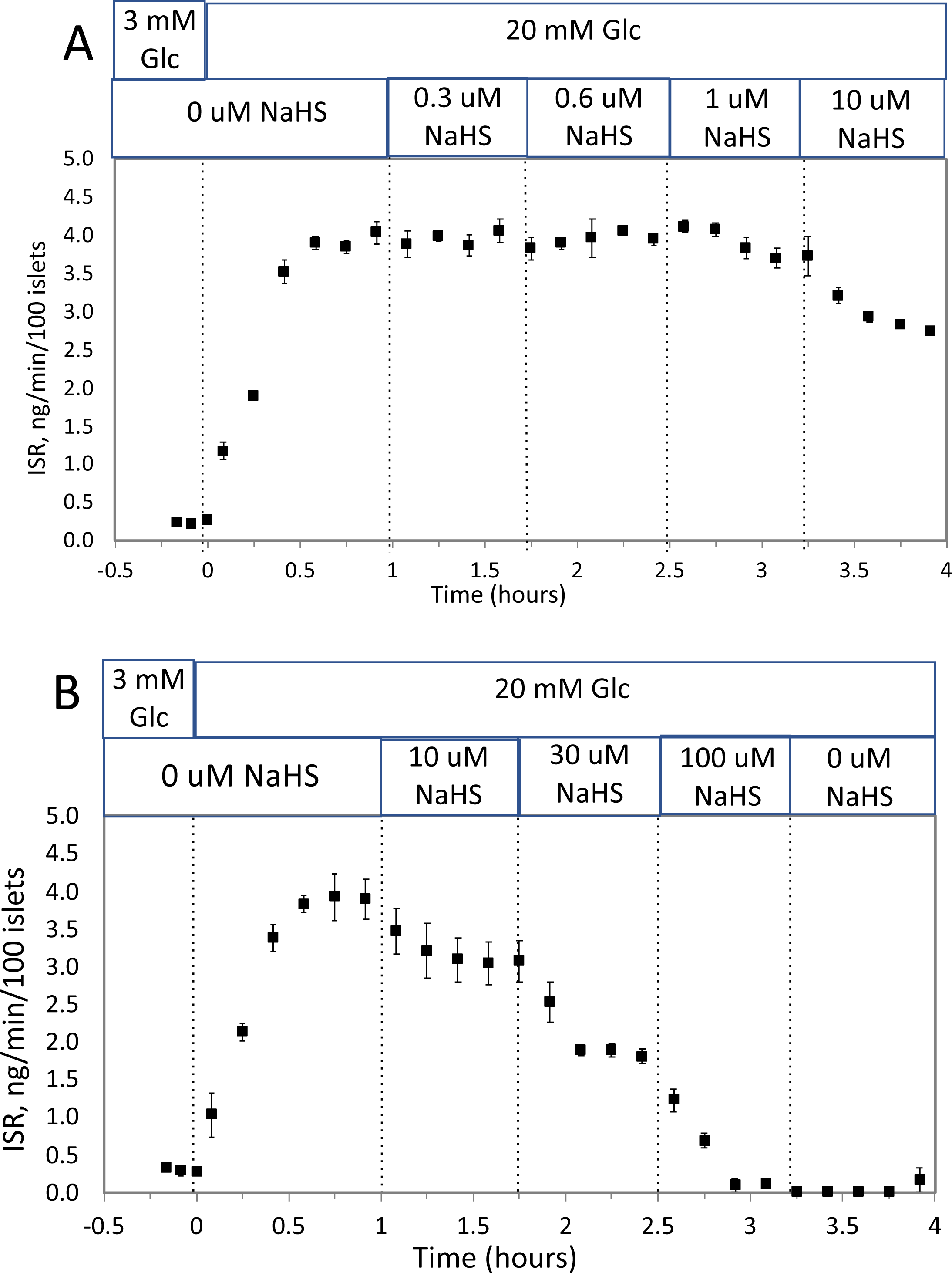
Effect of NaHS on ISR by islets. Rat islets (50/channel) were perifused (flow rate = 200 uL/min), and ISR was measured in response to glucose and exposure to incrementally increasing concentrations of aqueous NaHS as indicated. Data is average +/− SE, n =2.

**APPENDIX Fig. 2.**
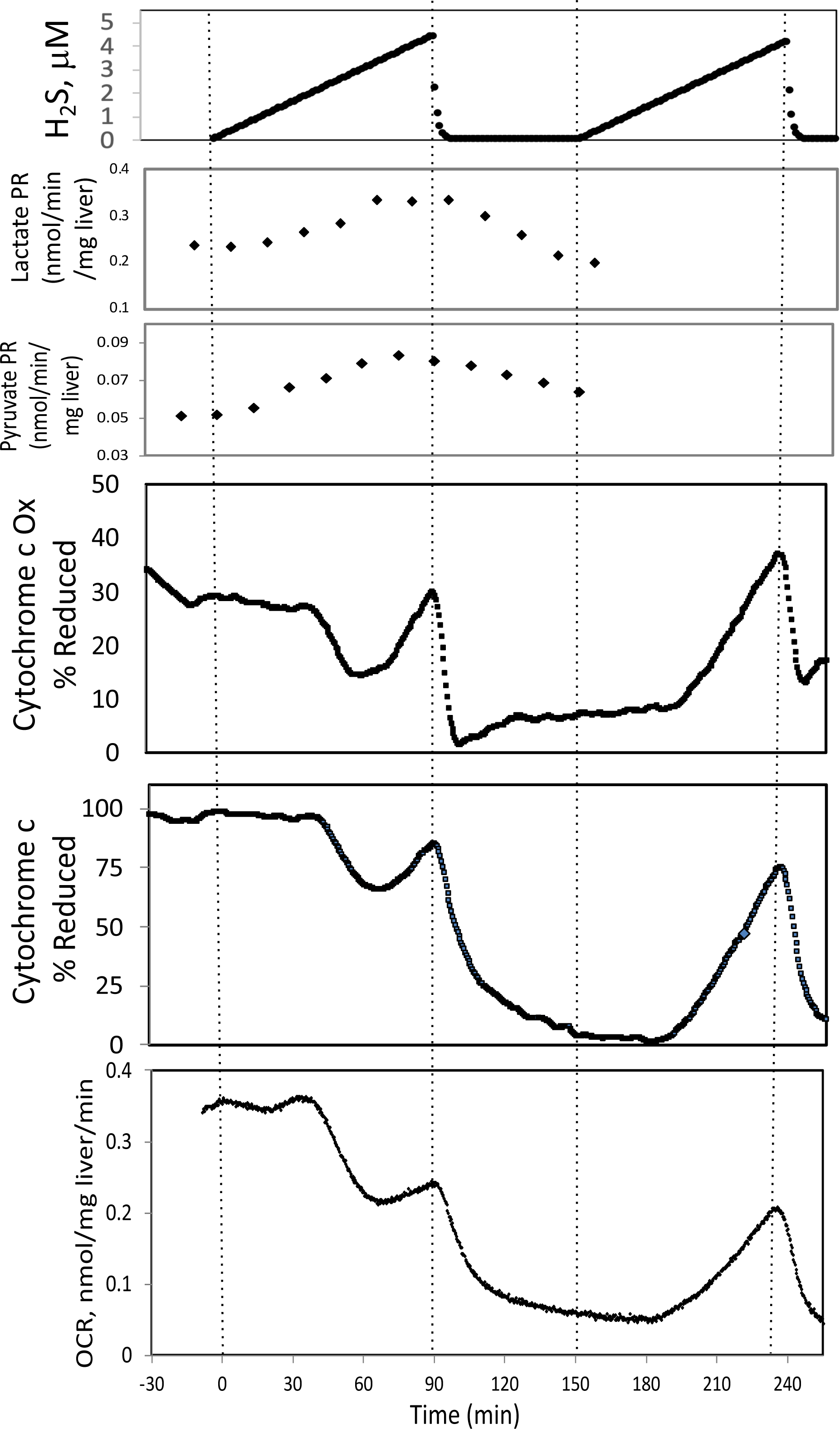
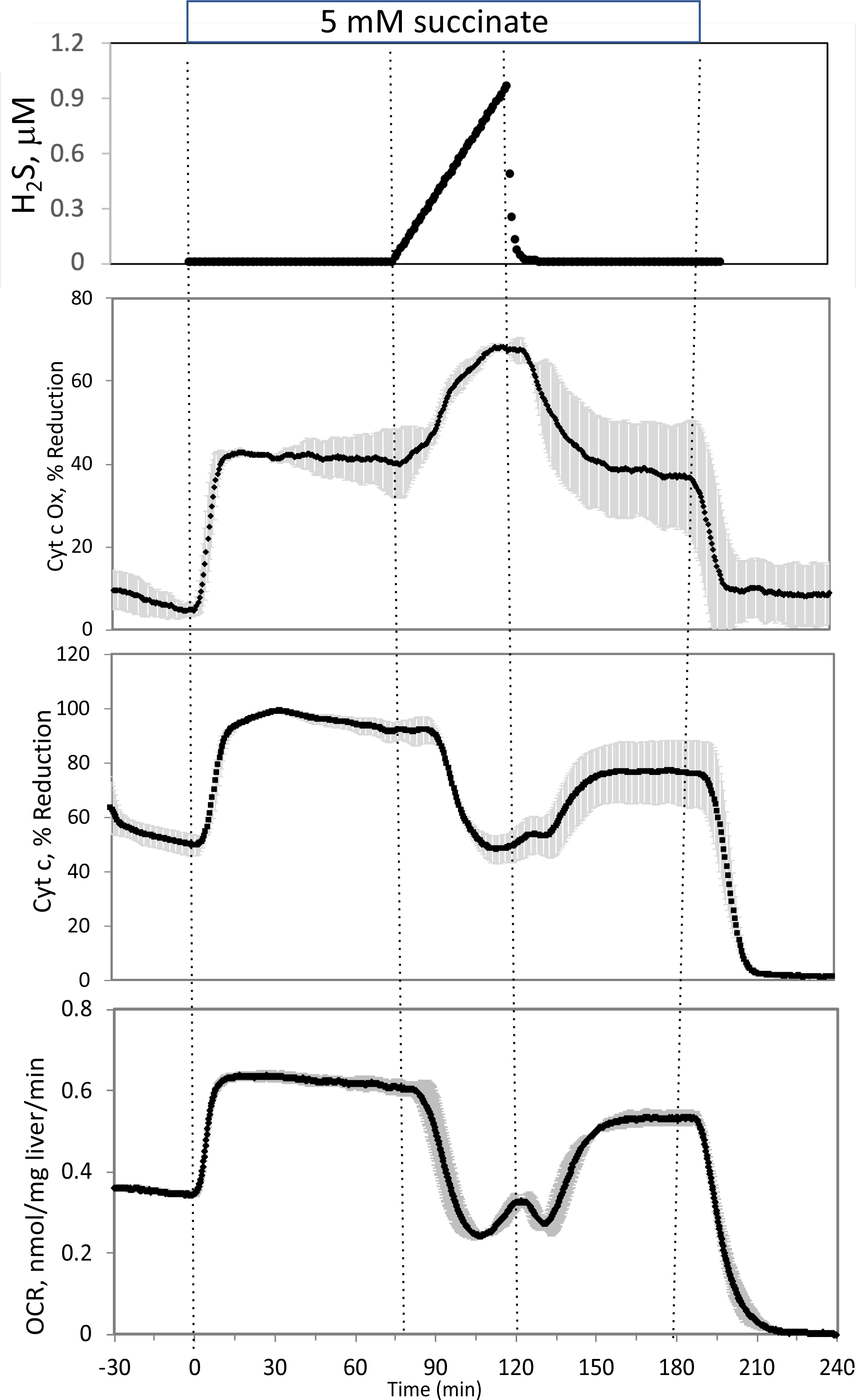
Effect of H_2_S on ETC and lactate/pyruvate in rat liver. In the absence (A, n=1)) or presence of 5 mM succinate (B, n=2), liver slices were exposed to increasing levels of H_2_S until the gas was purged as indicated. OCR, reduced cytochrome c and cytochrome c oxidase were measured in real time. Fractions were collected for subsequent measurement of lactate and pyruvate in (A) only.

